# Chronic Gq activation of ventral hippocampal neurons and astrocytes differentially affects memory and behavior

**DOI:** 10.1101/2022.09.15.508157

**Authors:** Rebecca L. Suthard, Alexandra L. Jellinger, Michelle Surets, Monika Shpokayte, Angela Y. Pyo, Michelle D. Buzharsky, Ryan A. Senne, Kaitlyn Dorst, Heloise Leblanc, Steve Ramirez

## Abstract

Network dysfunction is implicated in numerous diseases and psychiatric disorders, and the hippocampus serves as a common origin for these abnormalities. To test the hypothesis that chronic modulation of neurons and astrocytes induces impairments in cognition, we activated the hM3D(Gq) pathway in CaMKII+ neurons or GFAP+ astrocytes within the ventral hippocampus across 3, 6 and 9 months. CaMKII-hM3Dq activation impaired fear extinction at 3 months and acquisition at 9 months. Both CaMKII-hM3Dq manipulation and aging had differential effects on anxiety and social interaction. GFAP-hM3Dq activation impacted fear memory at 6 and 9 months. GFAP-hM3Dq activation impacted anxiety in the open field only at the earliest time point. CaMKII-hM3Dq activation modified the number of microglia, while GFAP-hM3Dq activation impacted microglial morphological characteristics, but neither affected these measures in astrocytes. Overall, our study elucidates how distinct cell types can modify behavior through network dysfunction, while adding a more direct role for glia in modulating behavior.

**Highlights:**

- CaMKII- and GFAP-Gq activation impacted memory, anxiety, and social behaviors.
- Novel environment exploration was affected by CaMKII- and GFAP-Gq activation.
- CaMKII-Gq modified microglial number, while GFAP-Gq affected microglial morphology.
- Neither cell manipulation affected astrocytic number or morphology.

## 1) Introduction

Cellular disturbances in the inhibitory and excitatory balance of brain circuits have been implicated in various neurodegenerative disorders such as Alzheimer’s Disease (AD), Multiple Sclerosis (MS) and Parkinson’s Disease (PD) (Lauterborn et al. 2021; Bi et al. 2020; Vico Varela et al. 2019; Lerdkrai et al. 2018; Campanelli et al. 2022; Ellvardt et al. 2018). Additionally, these imbalances, such as neuronal hyperactivity or hypoactivity, have been demonstrated to be core features of psychiatric disorders, including Post-Traumatic Stress Disorder (PTSD), Schizophrenia, and Major Depressive Disorder (MDD) (Badura-Brack et al. 2018; Fang et al. 2018; Clancy et al. 2017; Heckers & Konradi et al. 2014; Vadodaria et al. 2019; Helm et al. 2018). Interestingly, psychiatric disorders are often comorbid with neurodegenerative diseases, especially in AD and MDD (Martin-Sanchez et al. 2021). Along similar lines, psychiatric disorders such as MDD and PTSD significantly elevate the risk of developing neurodegenerative diseases such as AD (Flatt et al. 2018; Ownby et al, 2006; Byers & Yaffe, 2011). Thus, imbalances in circuit-level activity provide a putative shared mechanism underlying different brain and may serve as a therapeutic target.

Recently, many studies have investigated the role of direct brain modulation for the treatment and prevention of disease (Johnson et al., 2013; Gauthier et al. 2022), though the identity of the perturbed cells remains relatively unknown. For instance, stimulation of brain cells via transmagnetic stimulation (TMS), which is an approved therapy for treatment-resistant MDD, has promising therapeutic value when applied to AD, PD, stroke, schizophrenia, and MS (Weiler et al. 2020; Brys et al. 2016; van Lieshout et al. 2020; Cole et al. 2015; Nasios et al. 2018). However, understanding how to restore brain patterns through a cell-type specific and targeted manner remains elusive, especially when used for the treatment of brain diseases. Hyperactivity is a hallmark of many disorders and often occurs in areas of the brain involved in learning and memory. For instance, the hippocampus (HPC) is a deep brain structure crucially involved in learning and memory, and its connecting subregions are crucial for the formation and retrieval of episodic memories in particular (Eichenbaum, Otto & Cohen, 1992; Moser & Moser, 1998; Burgess, Maguire & O’Keefe, 2002; Eichenbaum & Fortin, 2003). Various clinical and preclinical studies have shown that in early disease states of AD, the HPC becomes hyperactive and, therefore, is a prime location for network dysfunction and disconnection from other brain regions (Palop et al. 2007; Busche et al. 2008; Busche et al. 2012; Palop & Mucke, 2016; Klink et al 2021; Anastacio, Matosi, & Ooi, 2022; Kuchibhotla et. al., 2009). One of the major subregions of the HPC that is directly affected in such diseases is along the ventral axis, especially ventral CA1 (vCA1; Maruszak & Thuret, 2014). Notably, the ventral HPC (vHPC) has direct connectivity to many downstream regions such as the basolateral amygdala (BLA), nucleus accumbens (NAc), medial prefrontal cortex (mPFC), and the hypothalamus, thus underscoring its role in modulating key aspects of memory and behavior (Fanselow & Dong, 2010; Gergues et. al., 2020; Ciocchi et. al., 2015; Padilla-Coreano et. al., 2016; Jimenez et. al., 2018; Zhou et. al., 2019; Phillips et. al., 2019). These connections have been specifically linked to processing the emotionally valenced components of episodic memories (Fanselow & Dong, 2010). Fittingly, dysfunction in a major hub like the vHPC may lead to deficiencies in numerous cognitive functions including learning, memory, emotional processing, reward-seeking behaviors, and stress responses. Thus, we sought to modulate vCA1 processing by chronically modulating its activity, to test the hypothesis that prolonged network perturbation would lead to cellular and behavioral abnormalities that may be unique to each cell type.

Within the vHPC, a number of studies have focused specifically on neuronal functioning and characterizing the roles of neurons in social, anxiety, goal-directed and fear-related behaviors (Jimenez et. al., 2018, Okuyama et. al., 2016; Jimenez et. al., 2020; Padilla-Coreano et. al., 2016; Ciocchi et. al., 2015; Fanselow & Dong, 2010). Notably, glial cells such as astrocytes, have remained understudied in this ventral axis of the HPC. Astrocytes have primarily been studied as support cells in the brain, aiding in neuronal metabolism, supporting the blood-brain barrier and providing homeostatic regulation of their local environment (Dehouck et. al., 1990; Simard & Nedergard, 2004; Kofuji & Newman, 2004; Paulson & Newman, 1987; Tsacopoulous & Magistretti, 1996; Wallraff et. al., 2006; Rouach et. al., 2008; Figley et. al., 2011). Their active role in synaptic regulation has now been fully recognized, as knowledge of their expression of receptors, transporters and bidirectional communication with neurons has grown (Porter et. al., 1997; Perea & Araque, 2005; Covelo & Araque, 2018; Araque et. al., 2001; Di Castro et. al., 2011; Haydon et. al., 2001; Bezzi & Volterra, 2001). It is now generally thought that astrocytes dynamically respond and modulate circuit activity via calcium-dependent and -independent release of gliotransmitters, such as glutamate, D-serine, and ATP, at the tripartite synapse (Araque et. al., 1999; Perea et. al., 2009; Parpura et. al., 1994; Koizumi et. al., 2005; Fellin et. al., 2004). As astrocytes play a pivotal role in many brain functions, modulation of their structure or activity has a major impact on cognition and behavior (Adamsky et. al., 2018; Kol et. al., 2020; Li et. al., 2020; Mederos et. al., 2021; Martin-Fernandez et. al., 2017; Skucas et. al., 2011). Recent work has shown that astrocytic calcium dysregulation in the cortex may be involved in the early stages of neurodegeneration and influence the onset of neuronal hyperactivity even before amyloid deposition in a mouse model of AD (Shah et. al., 2022). Astrocytes and neurons are heterogenous in their structure and function within and across brain regions, suggesting that perturbation of either cell type may elicit unique behavioral and cellular responses even within the same brain region (Oberheim, Goldman & Nedergaard, 2012; Matyash & Kettenmann, 2010; Buosi et. al., 2017; Zhang & Barres, 2010).

Here, we sought to study the differential effects of chemogenetic activation of the hM3D(Gq) pathway in excitatory neurons (calcium/calmodulin-dependent protein kinase II; CaMKII+) or astrocytes (glial fibrillary acidic protein; GFAP+) in the vHPC, as most studies have identified network hyper- or hypoactivity as a result of disease rather than how onset of this activity may produce pathological brain functioning. In this study, we tested the hypothesis that artificial chronic induction of network hyperactivity in both neuronal and astrocytic groups is sufficient to induce impairments in cognition and behavior, as well as changes in histological markers of cellular stress and inflammation. Our results demonstrate that chronic Gq pathway activation of CaMKII+ neurons or GFAP+ astrocytes in vHPC across multiple time points is sufficient to induce changes in behavior and histological markers. Behaviorally, CaMKII-Gq activation decreased contextual fear acquisition at the 9 month time point and increased fear during extinction at the 3 month time point. GFAP-Gq activation was generally impacted by our manipulation during CFC, recall and extinction at the 6 and 9 month time points. For anxiety-related tasks, CaMKII-Gq activation displayed a combinatory effect with aging to impact behavior. GFAP-Gq activation decreased anxiety-related behaviors at the 3 month time point in the open field, but only showed impacts of aging in the zero maze. For social interaction, there was a decrease in engagement with the social target in the CaMKII-Gq mice at 6 months. For GFAP-Gq mice, we observed only effects of aging on locomotion and social target engagement in the social interaction task. For novel environment exploration, there was an impact of chronic activation and aging on neuronal and astrocytic groups across all time points. Histologically, CaMKII-Gq mice displayed changes in microglial, but not astrocytic cell number in vHPC, while GFAP-Gq mice showed no differences in glial cell number. For morphology, only GFAP-hM3Dq activation impacted microglial characteristics, but not astrocytic, and CaMKII-hM3Dq activation did not affect either cell type. Our study thus provides a mechanistic approach to study how these cell types can be longitudinally engaged to produce behavioral deficits, which provides a putative link to disorders that have similar characteristic network dysfunctions (e.g. hyper- or hypoactivity), while also adding a more direct role for glia in modulating behavior.

## 2) Methods

### 2.1) Subjects

Wild-type male C57BL/6J mice (P29-35; weight 17-19g; Charles River Labs) were housed in groups of 2-5 mice per cage. The animal facilities (vivarium and behavioral testing rooms) were maintained on a 12:12-hour light cycle (0700-1900). Mice received food and water *ad libitum* before surgery. Following surgery, mice were group-housed with littermates and allowed to recover for a minimum of 4 weeks before experimentation to allow for viral expression. All subjects were treated in accord with protocol 201800579 approved by the Institutional Animal Care and Use Committee (IACUC) at Boston University.

### 2.2) Stereotaxic surgeries

For all surgeries, mice were initially anesthetized with 3.0% isoflurane inhalation during induction and maintained at 1-2% isoflurane inhalation through stereotaxic nose-cone delivery (oxygen 1L/min). Ophthalmic ointment was applied to the eyes to provide adequate lubrication and prevent corneal desiccation. The hair on the scalp above the surgical site was removed using Veet hair removal cream and subsequently cleaned with alternating applications of betadine solution and 70% ethanol. 2.0% lidocaine hydrochloride was injected subcutaneously as local analgesia prior to midsagittal incision of the scalp skin to expose the skull. 0.1mg/kg (5mg/kg) subcutaneous (SQ) dose of meloxicam was administered at the beginning of surgery. All animals received bilateral craniotomies with a 0.5-0.6 mm drill-bit for vCA1 injections. A 10uL Hamilton syringe with an attached 33-gauge beveled needle was slowly lowered to the coordinates of vCA1: −3.16 anteroposterior (AP), ± 3.10 mediolateral (ML) and both - 4.25/-4.50 dorsoventral (DV) to cover the vertical axis of vHPC. All coordinates are given relative to bregma (mm). A volume of 150nL (100nL/min) of AAV9-CaMKII-hM3Dq-mCherry, AAV9-CaMKII-mCherry, AAV5-GFAP-hM3Dq-mCherry or AAV5-GFAP-hM3Dq-mCherry was bilaterally injected into the vCA1 at both DV coordinates listed above. The needle remained at the target site for 7 minutes post-injection before removal. Incisions were sutured closed using 4/0 Non-absorbable Nylon Monofilament Suture [Brosan]. Following surgery, mice were injected with a 0.1mg/kg intraperitoneal (IP) dose of buprenorphine. They were placed in a recovery cage with a heating pad until fully recovered from anesthesia. Histological assessment verified bilateral viral targeting and data from off-target injections were not included in analyses.

### 2.3) Immunohistochemistry

Mice were overdosed with 3% isoflurane and perfused transcardially with cold (4°C) phosphate-buffered saline (PBS) followed by 4% paraformaldehyde (PFA) in PBS. Brains were extracted and kept in PFA at 4°C for 24-48 hours and transferred to a 30% sucrose in PBS solution. Long-term storage of the brains consisted of transferring to a 0.01% sodium azide in PBS solution until slicing. Brains were sectioned into 50 um thick coronal or sagittal sections with a vibratome and collected in cold PBS or 0.01% sodium azide in PBS for long-term storage.

Sections underwent three washes in PBS for 5-10 minutes each to remove the 0.01% sodium azide that they were stored in. Sections were washed three times for 5 minutes each in 0.2% Triton X-100 in PBS (PBST). Sections were blocked for 2 hours at room temperature (RT) in 1% bovine serum albumin (BSA) and 0.2%Triton X-100 in PBS on a shaker. Sections were incubated in primary antibody (1:1000 mouse anti-GFAP [NeuroMab]; 1:1000 rabbit polyclonal anti-Iba1/AIF1 [SySy]; 1:500 polyclonal guinea anti-NeuN/Fox3 [SySy], 1:1000 rabbit polyclonal anti-cFos [SySy],] 1:1000 rabbit polyclonal anti-RFP [Rockland]) made in the same 1% BSA/PBS/Triton X-100 solution at 4°C for 24-48 hours. Sections then underwent three washes for 5 minutes each in 0.2% PBST. Sections were incubated in secondary antibodies (1:1000 Alexa Fluor 488 anti-rabbit IgG [Invitrogen]; 1:1000 Alexa Fluor 488 anti-mouse IgG [Invitrogen]; 1:1000 Alexa Fluor 488 anti-guinea IgG [Invitrogen]) for 2 hours at RT. Sections then underwent three more washes in PBST. Sections were then mounted onto microscope slides (VMR International, LLC). Vectashield HardSet Mounting Medium with DAPI (Vector Laboratories, Inc) was applied and slides were coverslipped and allowed to dry for 24 hours at RT. Once dry, slides were sealed with clear nail polish around the edges and stored in a slide box in 4°C. If not mounted immediately, sections were stored in PBS at 4°C.

### 2.4) Behavioral assays

All behavior assays were conducted during the light cycle of the day (0700-1900) on animals at the 3, 6 and 9 month timepoints. Mice were handled for 3–5 days before all behavioral experiments began. Behavioral assays include open field, zero maze, social interaction, y-maze, contextual fear conditioning, recall and extinction tests. Before each day of behavioral testing, DCZ water was removed 1-2 hours from the home cage to allow for a decline in ligand concentration in the brain. DCZ administered I.P. in mice has been shown to peak 5-10 minutes after injection, declining to baseline around 120-150 minutes in mice at a low dose (Nagai et. al., 2021). Relatedly, there are pharmacokinetic differences with the P.O. route that have been shown in macaques, with serum DCZ concentrations more slowly increasing, but peaking at comparable or even higher levels than observed in the I.P. route (Mimura et. al., 2021). We chose to administer our ligand with the water-soluble administration route at a low dose of 3ug/kg to avoid any potential excitotoxicity resulting through stimulating cells for up to 9 months, and to mimic a more sustained dysregulation that may accumulate in severity over time.

#### Open field test

An open 61cm x 61cm arena with black plastic walls was used for the open field test, with a red-taped area of 45 cm x 45 cm in the middle delineating “center” from “edges”. A camera was placed above the open field in order to record video of the session. Mice were individually placed into the center of the chamber, and allowed to explore freely for 10 min. At the end of the session, each mouse was placed into a separate cage until all of its cage mates had also gone through the behavioral test. They were all placed back into the home cage once this occurred. An automated video-tracking system, AnyMaze, was used to measure total distance traveled, mean speed, number of entries into the center, and center mean visit time and total time spent in the center.

#### Zero maze test

The zero maze test is a pharmacologically-validated assay of anxiety in animal models that is based on the natural aversion of mice to elevated, open areas. It is composed of a 44cm wide ring with an outer diameter of 62cm. It contained four equal zones of walled (closed) and unwalled (open) areas. The entire ring is 8cm in height and sits at an elevation of 66cm off the ground. All animals at each timepoint were tested on the same day. Mice were placed in the closed area at the start of a 10 minute session. The following parameters were analyzed: distance traveled, mean speed, time spent in the open area, number of entries into the open area (head and body) and open area mean visit time. The maze was cleaned with 70% ethanol between mice. During the behavioral testing, the lighting levels remained constant and there were no shadows over the surface of the maze.

#### Social interaction test

An open area (60.96cm x 60.96cmin) with black walls was used for the social interaction test. Two inverted wire cups of diameter (10.16cm) and height (10.16cm) were placed in the arena in opposite corners, each set (12.7cm) away from the internal corner of the arena. Black permanent marker was used to draw a circle (15.24cm) on the floor of the area around the outside of the wire cup to demarcate a diameter (2.54cm) larger than that of the cup. A 2-3 month old male conspecific was placed into one wire cup (mouse cup), while the other cup was left empty (empty cup). Each test animal was placed into the middle of the arena and was allowed to freely explore for 10 minutes. A camera was mounted above the arena to record video of behavior for subsequent scoring using AnyMaze. The metrics scored included the total amount of time and number of entries for each region (mouse cup/social target, empty cup). Additionally, total distance traveled was scored.

#### Y-maze test

The Y-maze is a hippocampal-dependent spatial working memory task that requires mice to use external cues to navigate the identical internal arms. The apparatus consisted of a clear Plexiglass maze with three arms (36.5cm length, 7.5cm width, 12.5cm height) that were intersected at 120 degrees. A mouse was placed at the end of one arm and allowed to move freely through the maze for 10 minutes without any reinforcements, such as food or water. Entries into all arms were noted (center of the body must cross into the arm for a valid entry) and a spontaneous alternation was counted if an animal entered three different arms consecutively without repeat (e.g. ABC, CBA). Percentage of spontaneous alternation was calculated according to the following formula: [(number of alternations)/(total number of arm entries - 1) x 100. To prevent bias in data analysis, the test was carried out in a blind manner by the experimenter and behavior was analyzed blindly in AnyMaze.

#### Fear conditioning, recall and extinction

##### Fear Conditioning

Fear conditioning for all timepoints took place in mouse conditioning chambers (18.5 x 18.5 x 21.5cm) with metal-panel side walls, plexiglass front and rear walls and a stainless-steel grid floor composed of 16 grid bars (context A). The grid floor was connected to a precision animal shocker set to deliver foot shocks over a time span of 6 minutes or 360s with 1.5 mA 2 second shocks at 120s, 180s, 240s, 300s. A video camera was mounted to the ceiling of the chamber to record activity and fed into a computer running FreezeFrame software (Actimetrics). This software controlled stimulus presentations and recorded videos from two to four chambers simultaneously. The program determined movement as changes in pixel luminance over a set period of time. Freezing was defined as a bout of 1.25 s or longer without changes in pixel luminance and verified by an experimenter blind to treatment groups. The chambers were cleaned with 70% ethanol solution prior to each animal placement.

##### Recall

For the recall test, animals were placed into context A for a duration of 5 minutes or 300s 24 hours after fear conditioning.

##### Extinction

Contextual extinction for all time points took place in context A for 30 minutes or 1800s per session. Three extinction sessions were administered across three consecutive days beginning 24 hours after recall.

### 2.5) DCZ administration

#### Acute administration

A separate cohort of mice were subjected to the same surgical protocol described above. After waiting 4 weeks post-surgery for viral expression, mice were injected intraperitoneally (i.p.) with deschloroclozapine (DCZ). DCZ was mixed into solution with dimethyl sulfoxide (DMSO) and diluted in sterile saline for a final concentration of 30ug/mL administered at 0.3mg/kg. 90 minutes after injection, mice were time perfused to capture peak cFos protein levels, as described in recent literature documenting cFos levels in DCZ administered rats (Nentwig et. al., 2022). This injection protocol was performed with 3 mice per group [CaMKII-hM3Dq, CaMKII-mCherry, GFAP-hM3Dq and GFAP-mCherry] for histological assessment of cFos changes with acute administration of non-water soluble DCZ. It should be noted that mice in the CaMKII-hM3Dq group were the only subjects that appeared behaviorally to experience seizure activity. This seizure activity was not observed in any of the other groups during acute administration, nor during our chronic water-soluble administration over the course of 3, 6 and 9 month time points. We speculate that this is due to acute I.P. compared to P.O. dosing pharmacokinetics with rapid onset, high affinity and specificity in the hippocampus, a region prone to seizure activity.

#### Chronic administration

All animals had their home cage water supply replaced with water-soluble DCZ in distilled water (diH2O) every 5 days. Administration was calculated based on each mouse drinking approximately 6mL of water per day. A stock solution was created of 1-2mg of water soluble DCZ into 1 mL of DMSO and vortexed until in solution. 17ul of stock solution was combined with 1L of diH20 for a final concentration of 3ug/kg/day per mouse.

### 2.6) Imaging and cell counting

All coronal and sagittal brain slices were imaged through a Zeiss LSM 800 epifluorescence microscope with a 10x/20x/40x/64x objective using the Zen2.3 software.

Full slice images of RFP amplified hM3Dq expression (Figure 1B,D) were captured along with zoomed in a 300 x 300 tile (micrometer) z-stack at 20x magnification. 3-4 different slices were imaged for each animal for averaging, for each brain region of interest. Images of NeuN, GFAP, and Iba-1 were captured in a 300 x 300 single-tile (micrometers) z-stack at 20x magnification. 3-4 mice per group, with 18 single-tile ROIs covering the vHPC were used for cell counts of NeuN, GFAP and Iba-1. Morphological analysis of microglia (Iba1) and astrocytes (GFAP) was performed using 3DMorph, a MATLAB-based tool that analyzes glial morphology from 3-dimensional imaging data (York et. al., 2018). For cFos counts, 3 mice per group with 4 images (1182 x 1756 micrometers) each covering the vCA1 pyramidal cell layer 20x magnification were used. Each pyramidal cell layer ROI was hand-traced in ImageJ and cFos counts were normalized to the number of DAPI (%cFos/DAPI). NeuN, GFAP, Iba-1, DAPI and cFos counts were performed using Ilastik, a machine-learning-based image analysis tool (Berg et. al., 2019).

**Figure 1.**
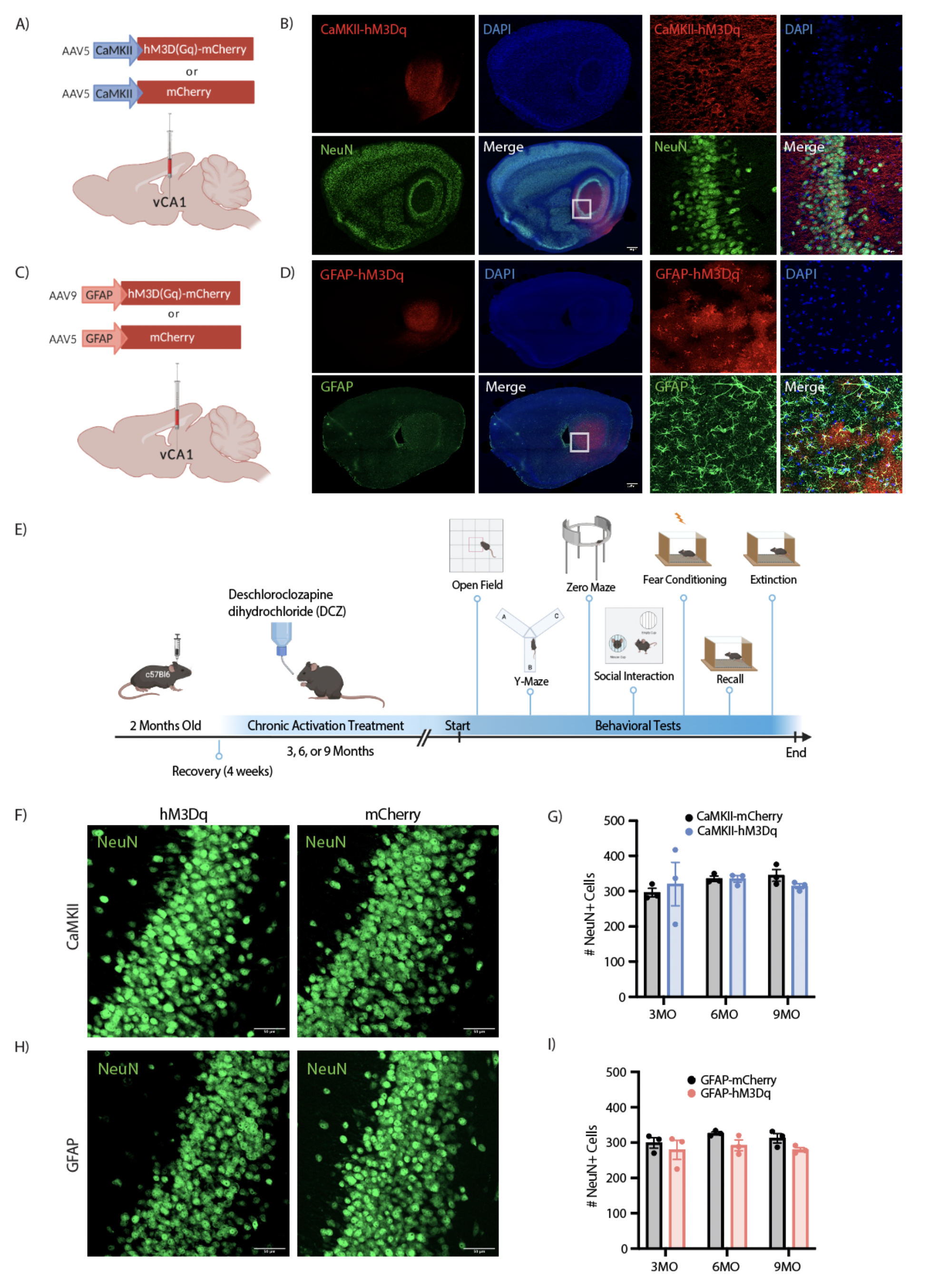
Chronic activation of Gq pathways in CaMKII+ neurons and GFAP+ astrocytes in vCA1. (A) Viral strategy schematic for the neuronal groups. The AAV5-CaMKII-hM3D(Gq)-mCherry or control vector AAV5-CaMKII-mCherry was bilaterally injected into the vCA1 region of wild type mice. (B) Representative images of RFP/NeuN+ costaining (red and green, respectively) and DAPI+ cells (blue). (C) Viral strategy schematic for the astrocyte groups. The AAV9-GFAP-hM3D(Gq)-mCherry or control vector AAV5-GFAP-mCherry was bilaterally injected into the vCA1 region of wild type mice. (D) Representative images of RFP/GFAP+ co-staining (red and green, respectively) and DAPI+ cells (blue). (E) Schematic representation of chronically activating neuron or astrocyte Gq receptors through administration of the water-soluble DREADD ligand deschloroclozapine dihydrochloride (DCZ) for either 3, 6, or 9 months. Mice underwent a battery of behavioral tests at each end point (F-I) Histological assessment of NeuN+ cells to determine whether our manipulations were killing vCA1 neurons in the CaMKII groups (F and G) and/or GFAP groups (H and J). NeuN counts were assessed with a two-way analysis of variance (ANOVA) with time point and group as factors. Error bars indicate SEM. p ≤ 0.05, **p ≤ 0.01, ***p ≤ 0.001, ****p ≤ 0.0001, ns = not significant. Per group: n= 3 mice x 18 tiles (region of interest (ROI): vCA1) each were quantified for statistical analysis of NeuN counts. Scale bars indicate 50 micrometers.

For Gq receptor fluorescence intensity measures (Supplemental Figure 7), sagittal slices of vHPC for CaMKII- and GFAP-hM3Dq groups were imaged for mCherry expression, (no immunohistochemistry was used to amplify the signal). Here, we imaged DAPI and Gq-mCherry expression for 3 mice with 3 vHPC ROI’s each (2938 x 3513 micrometers), across groups (GFAP/CaMKII-hM3Dq) x time points (3,6,9 months). These images were taken using a 10x objective over the entire vHPC, with the ROI maintained across images, slices and animals. All imaging parameters were identically maintained across all animals and slices for fluorescence intensity comparison. Fluorescence intensity measures were conducted in ImageJ using two methods to ensure consistency. Method 1: select vHPC ROI (entire image taken), Analyze, Set Measurements, Select ‘mean gray value’, Analyze, Measure. The mean gray value was used as our measure, as all images were the same area and all settings were maintained, so this should be standardized across all mice and groups, unlike ‘integrated density’ that takes different areas into account. Method 2: select 3 ROIs in the fluorescent region and 3 ROIs in the dark ‘background’ of each image at random, then use the discussed steps to gather ‘mean gray value’, ‘area’ and ‘integrated density’ values for each small ROI. These values were averaged for each image, then averaged again for each mouse for a final ‘background’ and ‘fluorescent’ mean gray, area and integrated density value. Using these metrics, we calculated the average corrected total cell fluorescence (CTCF) using integrated density of the ‘fluorescent’ region - (area of the ‘fluorescent’ ROI x mean gray fluorescence of ‘background’ region).

### 2.7) Statistical Methods

Data was analyzed using GraphPad Prism. All data were tested for normality using Shapiro-Wilk and Kolmogorov–Smirnov tests. Outliers were removed prior to statistical analysis using the ROUT method recommended by GraphPad Prism, which uses identification from nonlinear regression. We chose a ROUT coefficient Q value of 10%(False Discovery Rate), making the threshold for outliers less-strict and allowing for an increase in power for outlier detection. To analyze differences between groups and across timepoints, we used: Two-way ANOVAs (between-subject factor: Group; within-subject factor: Timepoint). Alpha was set to p<0.05. Post-hoc analyses were run using Tukey’s-multiple comparisons test with a 95% CI. To analyze differences between groups and across time within a single session we used: Two-way repeated measures (RM) ANOVAs (between-subject factor: Group; within-subject factor: Time). Alpha level was set to 0.05. Post-hoc analyses were run using Šídák’s multiple comparisons test with a 95% CI. To analyze differences between groups, we used: Independent t-tests [CaMKII-hM3Dq vs. mCherry; GFAP-hM3Dq vs. mCherry]. Alpha was set to 0.05.

### 2.8) Reagents and Resources

|  | SOURCE | IDENTIFIER |
| --- | --- | --- |
| Experimental Models: Organisms/Strains |  |  |
| Wild-type male C57Bl/6 | Charles River Labs | C57BL/6 |
| **Antibodies** |  |  |
| Rabbit polyclonal anti-Iba1/AIF1 | SySy | 234 003 (catalog #) |
| Mouse monoclonal anti-GFAP | NeuroMab | 75-240 (catalog #) |
| Guinea polyclonal anti-NeuN/Fox3 | SySy | 266-004 (catalog #) |
| Rabbit polyclonal anti-cFos | SySy | 226-008 (catalog #) |
| Rabbit polyclonal anti-RFP | Rockland | 600-401-379-RTU |
| Goat anti rabbit Alexa Fluor 488 Secondary antibody | Invitrogen | A-11008 (catalog #) |
| Goat anti-guinea pig Alexa Fluor 488 Secondary antibody | Invitrogen | A-11073 (catalog #) |
| Goat anti-mouse Alexa Fluor 488 Secondary antibody | Invitrogen | A-11001 (catalog #) |
| Goat anti-rabbit Alexa Fluor 555 Secondary antibody | Invitrogen | A27039 (catalog #) |
| **Chemicals** |  |  |
| Triton X-100 | FisherScientific | NC0478124 |
| Paraformaldehyde (PFA) | Sigma-Aldrich | 158127-3KG |
| Bovine serum albumin (BSA) | Sigma-Aldrich | B4287-25G |
| Deschloroclozapine | HelloBio | HB8555 |
| Deschloroclozapine dihydrochloride | HelloBio | HB9126 |
| Vectashield Hard Set Mounting Medium with DAPI | Vector Laboratories, Inc. | H-1500-10 |
| **Software/Programs** |  |  |
| Fiji (ImageJ) |  | https://fiji.sc/ |
| FreezeFrame4 | Coulbourn Instruments |  |
| AnyMaze | Stoelting Co. |  |
| Prism | GraphPad Software | v9.4.1 |
| Ilastik | Berg et. al., 2019 |  |
| 3DMorph | York et. al., 2018 |  |

| Viral Constructs |  |  |
| --- | --- | --- |
| AAV9-CaMKII-hM3Dq-mCherry | AddGene | 50476-AAV9 |
| AAV9-CaMKII-mCherry | AddGene | 114469-AAV9 |
| AAV5-GFAP-hM3Dq-mCherry | AddGene | 50487 -AAV5 |
| AAV5-GFAP-hM3Dq-mCherry | AddGene | 58909-AAV5 |

## 3) Results

### 3.1) Induction of network dysfunction in neurons or astrocytes across time

Network dysfunction is an early hallmark of neurodegenerative diseases such as Alzheimer’s Disease (AD) both in humans and rodent models (Lauterborn et. al., 2021; Vico Varela et. al., 2019). However, it is unknown whether chronic activation of astrocytes or neurons in the vHPC may induce behavioral and histological changes related to neurodegeneration in the brains of wild type mice. To address this question, we used chemogenetic methods to chronically activate the hM3D(Gq) pathway in either CaMKII+ neurons or GFAP+ astrocytes in the vCA1 region of the HPC. For the neuronal groups, wild type mice were injected bilaterally with AAV5-CaMKII-hM3D(Gq)-mCherry or control vector AAV5-CaMKII-mCherry to selectively express excitatory DREADD receptors in vCA1 (Figure 1A-B). For the astrocyte groups, C57BL/6 wild type mice were injected bilaterally with AAV9-GFAP-hM3D(Gq)-mCherry or control vector AAV5-GFAP-mCherry to selectively express DREADD receptors in astrocytes within vCA1 as well (Figure 1C-D). Our viruses were robustly expressed in each cell type, as indicated by co-staining with RFP/GFAP+ and RFP/NeuN+ for DREADD receptors and their unique cell markers (Figure 1B,D). After 4 weeks of recovery post-surgery, a water-soluble DREADD ligand, deschloroclozapine dihydrochloride (DCZ) was administered via the animal’s daily water to chronically activate the hM3D(Gq) receptors of neurons or astrocytes (Figure 1E) (Nagai et. al., 2020). DCZ was selected for its high potency, selectivity and affinity for hM3D(Gq) receptors compared to previously used DREADD agonists, such as clozapine-N-ozide (CNO) and compound 21 (C21)(Nagai et. al., 2021). Even at low doses (0.001-0.1 mg/kg), DCZ enhances neuronal activity when bound to hM3Dq receptors expressed in the mouse and monkey brain (Nagai et. al., 2021; Nentwig et. al., 2022). Chronic activation of these DREADD-receptors via the intraperitoneal (I.P.) route may cause pain or unnecessary stress for the animal, even with adequate habituation (Mimura et. al., 2021). To circumvent this issue, recent studies have employed administration of CNO with water or food (Padilla et. al., 2017; Zhan et. al., 2019; Fernandez et. al., 2018). DCZ has been used recently in this manner for voluntary oral (P.O.) administration, and these experiments have demonstrated no significant differences between the I.P. and P.O. routes on behavior at the same dose of 0.5mg/kg (Ferrari et. al., 2022). Limitations of this method include decreased control over timing and dose of administration (Mimura et. al., 2021), but this voluntary P.O. administration was deemed superior for chronic stimulation of these receptors for our experiments here. Given this, mice underwent chronic activation for 3, 6 or 9 months, for a total of 12 groups: 3 Month GFAP mCherry and 3 Month GFAP hM3Dq; 3 Month CaMKII mCherry and 3 Month CaMKII hM3Dq; 6 Month GFAP mCherry and 6 Month GFAP hM3Dq; 6 Month CaMKII mCherry and 6 Month CaMKII hM3Dq; 9 Month GFAP mCherry and 9 Month GFAP hM3Dq; 9 Month CaMKII mCherry and and 9 Month CaMKII hM3Dq (Figure 1E). At each group’s designated timepoint, mice were subjected to a battery of behavioral tasks including open field, zero maze, social interaction, y-maze, contextual fear conditioning (CFC), recall and extinction (Figure 1E). Brain tissue was obtained following behavioral testing and was used to perform various histological assessments. Finally, to measure whether our manipulations were killing neurons within the vCA1, a confounding variable, we quantified a neuron-specific nuclear marker, NeuN (Figure 1F-I; Supplemental Figure 3A-F). Analysis of CaMKII-hM3Dq and -mCherry groups revealed no interaction between time point and group for the number of NeuN+ cells, nor did each factor alone contribute to a change in NeuN+ cells (Figure 1F-G)(Two-way ANOVA; Interaction F(2,12)=0.5069, p=0.6147; Timepoint: F (2, 12) = 0.5594, p=0.5858; Group: F (1, 12) = 0.01321, p=0.9104). However, while analysis of the GFAP groups revealed no interaction between time point and group for the number of NeuN+ cells, there was an impact of group alone (Two-way ANOVA; Interaction: F (2, 12) = 0.1286, p=0.8805; Timepoint: F (2, 12) = 0.8680, p=0.4446; Group: F (1, 12) = 5.227, p=0.0412)(Figure 1H-I). This suggests that our manipulation of the GFAP-Gq pathway is only mildly affecting the number of NeuN+ cells in a time-independent manner, possibly indicating that there is an initial ‘insult’ or loss of neurons that does not worsen over time with our chronic Gq activation.

To confirm the successful acute activation of our Gq-DREADD receptors, we performed intraperitoneal injection of deschloroclozapine (DCZ) in separate cohorts of CaMKII-hM3Dq, CaMKII-mCherry, GFAP-hM3Dq and GFAP-mCherry mice and sacrificed them 90 minutes later to stain for peak cFos protein levels. Additionally, we sacrificed wild-type mice without the administration of DCZ to understand baseline cFos levels in vHPC and provide insight into the inherent change in cFos due to I.P. injection alone or with the addition of the ligand in the brain. Importantly, recent evidence has shown that DCZ and saline administration did not show any significant differences in cFos levels in rats lacking DREADD receptor expression, suggesting that DCZ alone does not impact cFos levels (Nentwig et. al., 2022). We found that across CaMKII-hM3Dq+DCZ, CaMKII-mCherry+DCZ, and -DCZ groups, there was a significant difference in mean cFos levels (One-way ANOVA; F(2,6)= 137.4, p<0.0001). Specifically, multiple comparisons revealed that DCZ administration significantly increased cFos levels between CaMKII-mCherry and CaMKII-hM3Dq + DCZ groups, indicating an increase in neuronal activity in vHPC due to presence of the ligand (Tukey’s: p<0.0001) (Supplemental Figure 6A-B, E-F). Additionally, the -DCZ control was not significantly different than the CaMKII-mCherry + DCZ group, suggesting that DCZ alone is not impacting cFos levels in the absence of active receptor (Tukey’s: p =0.6732)(Supplemental Figure 6A-B, E-F). This neuronal result is consistent with previous literature showing that hM3Dq activation of neurons increases neuronal activity at low doses of DCZ (Nagai et. al., 2021).

Additionally, GFAP-hM3Dq+DCZ, GFAP-mCherry+DCZ, and the -DCZ control group had significant differences in mean cFos levels (One-way ANOVA; F(2,6) = 5.907, p=0.0382). Specifically, *post-hoc* multiple comparisons revealed that GFAP-hM3Dq+DCZ trended towards a decrease in cFos levels in vCA1 of the HPC (Tukey’s: p=0.0578)(Supplemental Figure 6C-E, G). Additionally, it is notable that there was no significant pairwise difference between the GFAP-hM3Dq+DCZ and -DCZ control group (Tukey’s: p=0.9986)(Supplemental Figure 6C-E, G). But we observed a trend towards a difference in cFos levels across the GFAP-mCherry+DCZ and -DCZ control mice (Tukey’s: p=0.0543)(Supplemental Figure 6C-E, G). This result appears to suggest that DCZ administration in the GFAP-mCherry+DCZ group is driving a small increase in cFos levels and our GFAP-hM3Dq+DCZ is not being activated by the ligand, maintaining similar levels to ‘baseline’ -DCZ cFos. However, we speculate that the I.P. injection alone may be driving this apparent increase in the GFAP-mCherry+DCZ group as this region of the brain processes stress and negative emotional valence, while our -DCZ mice did not receive an I.P. injection. If this were the case, performing the same ‘baseline’ experiment with the addition of saline I.P. instead of -DCZ alone would likely show similar levels of cFos to the GFAP-mCherry+DCZ group. Thus, we would still observe a modest decrease in cFos in the GFAP-hM3Dq+DCZ compared to GFAP-mCherry+DCZ control group in our chronic experiment. The small decrease in cFos that we observe in the astrocytic hM3Dq group may seem counterintuitive, but it has become clear in recent studies that modulation of astrocytes and neurons with chemogenetics results in differential effects on cFos expression and behavior. For instance, Gi- and Gq-activation in astrocytes stimulates the release of extracellular glutamate and increases neuronal excitability, suggesting that inhibitory signaling may be a unique property to neurons (Durkee et. al., 2019). Another study has shown that Gq- and Gi-pathways in astrocytes drive synaptic potentiation in hippocampal CA1 via Ca2+-dependent and Ca2+-independent mechanisms, respectively (Van Den Herrewegen et. al., 2021). This suggests that our astrocytic Gq manipulation strategy may be inducing unique cellular outcomes compared to neuronal Gq activation, and future studies can seek to tease out the differential mechanisms underlying each.

There are clear limitations to only measuring cFos with acute administration of DCZ, as receptor expression over these long time points may diminish. To provide evidence that we are maintaining receptor expression, we quantified fluorescence intensity of hM3Dq receptor expression in vHPC across the experimental groups. We speculate that cFos levels measured at the chronic 3, 6 and 9 month time point may show similar cFos levels across groups, despite any changes in regional activity, as the brain may attempt to restore homeostasis with increases in cellular activity over time. This would render any cFos levels at each time point potentially inconclusive read-outs of continual activation of the DREADD receptors. Future experiments will be needed to confirm the electrophysiological responses in the vHPC across chronic manipulations to confirm at the most basic level any decline in receptor functionality. Overall, we show that our acute administration of the DCZ ligand activated the Gq pathway successfully and induced cellular changes in the brain area of interest with acute administration. Additionally, we show the maintenance of Gq-mCherry expression across the 3, 6 and 9 month time points to better understand if the DREADD receptor remains present and potentially active across long time periods (Supplemental Figure 7). Measurements for CaMKII-hM3Dq mice across the three time points did not reveal any significant differences in mean gray value or corrected total cell fluorescence (CTCF) across time ([Mean gray: One-way ANOVA; F (2, 6) = 0.6852, p=0.5395][CTCF: One-way ANOVA; F (2, 6) = 0.6155, p=0.5713)(Supplemental Figure 7A-C). For the GFAP-hM3Dq mice, we did not observe any differences in mean gray or CTCF across any time point ([Mean gray: One-way ANOVA; F (2, 6) = 0.7896m p=0.4961][CTCF: One-way ANOVA; F (2, 6) = 0.7541, p=0.5103])(Supplemental Figure 7D-F).

Together, these results provide evidence that chronic administration of DCZ does not produce differences in the number of NeuN+ cells in CaMKII-hM3Dq and mCherry groups, but there is an effect of our manipulation across the GFAP-hM3Dq and mCherry groups. Additionally, acute administration of DCZ results in differential cellular changes in vCA1, with an increase in cFos in the CaMKII-hM3Dq group and a decrease in cFos in the GFAP-hM3Dq group compared to their mCherry controls. Finally, maintenance of receptor expression across all time points provides evidence that there may be successful activation of DREADD receptors even at long time periods after surgery, but this is not a deterministic measure of success within the confines of our experiment.

### 3.2) Chronic Gq activation of CaMKII+ neurons impacts freezing behavior during contextual fear acquisition and maintenance

To test the hypothesis that chronic neuronal Gq activation is sufficient to induce behavioral deficits across all timepoints, we first subjected our mice to CFC, recall, and three days of extinction (Figure 1E). For average percent freezing during CFC, there was a significant interaction between time point [3 month v. 6 month v. 9 month] and group [CaMKII-hM3Dq v. -mCherry] (Two-way ANOVA; Interaction: F (2, 60) = 9.094, p=0.0004; Time point: F (2, 60) = 11.04, p<0.0001; Group: F (1, 60) = 9.496, p=0.0031)(Figure 2A; iv). Subsequent *post hoc* tests demonstrated that 9 month CaMKII-mCherry mice had significantly higher levels of freezing than 3 month CaMKII-mCherry mice (Tukey’s; p<0.0001), indicating an increase in fear with aging as shown in previous studies (Yanai & Endo, 2021)(Figure 2A; iv). Most notably, at the 9 month time point, CaMKII-hM3Dq mice had lower overall levels of freezing compared to CaMKII-mCherry controls during CFC (Tukey’s, p=0.0013)(Figure 2A; iv). Within the CFC session at 3 months, there was no significant interaction between time bins [seconds; 60, 120, 180, 240, 300, 360] and groups [CaMKII-hM3Dq vs. mCherry], but there was an individual effect of time bin alone. This is consistent with the natural change in freezing over the course of a CFC session as repeated shocks are administered (Two-way ANOVA RM; Interaction: F (5, 100) = 1.123, p=0.3532; Time: F (3.377, 67.55) = 71.55, p<0.0001; Group: F (1, 20) = 2.595, p=0.1229; Mouse: F (20, 100) = 3.404, p<0.0001)(Figure 2A; i). Within the CFC session at 6 months, there was no interaction between time bins and group, but there are individual effects of time bin and group alone. (Two-way ANOVA RM; Interaction: F (5, 110) = 1.529, p=0.1866; Time: F (3.017, 66.38) = 70.03, p<0.0001; Group: F (1, 22) = 5.968, p=0.0231; Subject: F (22, 110) = 2.581, p=0.0006)(Figure 2A; ii). There were differences in freezing dependent on time bins similar to the 3 month mice, as well as general differences in freezing levels in the CaMKII-mCherry compared to the CaMKII-hM3Dq experimental group. Finally, for 9 months, there was a significant interaction between time and group within session for CFC (Two-way ANOVA RM; Interaction: F (5, 90) = 2.484, p=0.0373; Time: F (3.214, 57.84) = 58.85, p<0.0001, Group: F (1, 18) = 15.93, p=0.0009, Mouse: F (18, 90) = 2.771, p=0.0008)(Figure 2A; iii). *Post hoc* multiple comparisons revealed that this was driven by lower levels of freezing in the CaMKII-hM3Dq group at the 180, 240 and 360 second time bins during CFC (Šidák multiple comparisons: 180: p<0.0001; 240: p=0.0044; 360: p=0.0168)(Figure 2A; iii).

**Figure 2.**
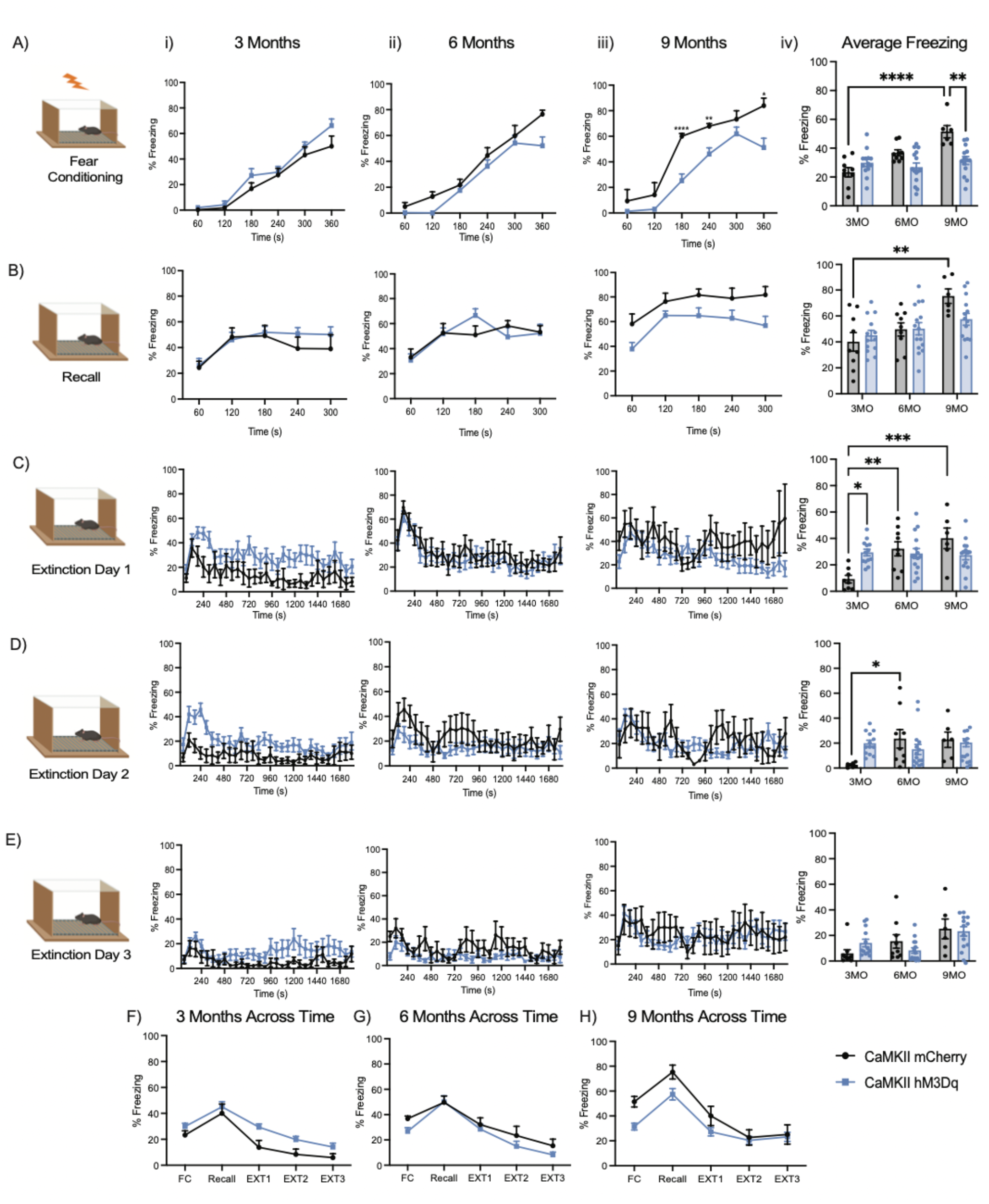
Chronic Gq activation of CaMKII+ neurons induces behavioral changes in contextual fear acquisition and maintenance. (A-E) Percent freezing levels and freezing percentage across time (seconds) of 3, 6, and 9 month time points. (A) CaMKII groups during contextual fear conditioning (CFC), where mice received four, 1.5mA foot shocks at 120, 180, 240 and 300 seconds during a 360 second session, (B) recall session for 360 seconds, where mice did not receive the aversive stimulus, (C) extinction day 1, (D) day 2 and (E) day 3, where mice received no foot shock for 1800 second sessions. (F-H) Average percent freezing (%) across fear conditioning, recall and extinction days 1-3 for CaMKII-hM3Dq and mCherry groups at 3 month (F), 6 month (G) and 9 month (H) time points. Average percent freezing was assessed with a two-way analysis of variance (ANOVA) with time point and group as factors. Freezing percentage across the 1-minute time bins [60, 120, 180, 240, 300, 360] was assessed with a two-way ANOVA with repeated measures (RM) with time (seconds) and group as factors. Average percent freezing across days [FC, Recall, EXT1, EXT2, EXT3] was assessed with a two-way ANOVA with repeated measures (RM) with day and group as factors. Tukey’s and Sidak’s post hoc tests were performed where applicable. Error bars indicate SEM. p ≤ 0.05, **p ≤ 0.01, ***p ≤ 0.001, ****p ≤ 0.0001, ns = not significant. 3 Month: hM3Dq (n=13), mCherry (n=9); 6 Month: hM3Dq (n=15), mCherry (n=9); 9 Month: hM3Dq (n=14), mCherry (n=6).

For contextual recall, there was no significant interaction between time point and group for average freezing, but time point alone had an individual effect, suggesting that aging alone is affecting freezing levels (Twoway ANOVA; Interaction: F (2, 60) = 2.327, p=0.1064; Timepoint: F (2, 60) = 9.508, p=0.0003; Group: F (1, 60) = 0.8738, p=0.3536)(Figure 2B; iv). Within recall sessions at 3 and 6 months, there were no significant interactions between time bin and group, but time bin alone had an effect on freezing levels (3 month: Two-way ANOVA RM; Interaction: F (4, 80) = 0.9819, p=0.4223; Time: F (3.089, 61.79) = 10.53, p<0.0001; Group: F (1, 20) = 0.4380, p=0.5157; Subject: F (20, 80) = 7.617, p<0.0001; 6 month: Two-way ANOVA RM; Interaction: F (4, 88) = 1.979, p=0.1046; Time: F (3.204, 70.49) = 10.38, p<0.0001; Group: F (1, 22) = 0.005898, p=0.9395; Subject: F (22, 88) = 6.368, p<0.0001)(Figure 2B; i-ii). Finally, at 9 months there was no interaction between time bin and group, but each factor alone, time and group, affected freezing levels within session (Two-way ANOVA RM; Interaction: F (4, 72) = 0.4759, p=0.7533; Time: F (3.048, 54.87) = 8.045, p=0.0001; Group: F (1, 18) = 5.111, p=0.0364; Subject: F (18, 72) = 5.851, p<0.0001) (Figure 2B; iii).

For extinction day 1, there was an interaction between time point and group for average percent freezing (Two-way ANOVA; Interaction: F (2,59) = 7.787, p=0.0010; Timepoint: F (2, 59) = 6.174, p=0.0037; Group: F (1,59) = 0.1405, p=0.7091)(Figure 2C; iv). Subsequent *post hoc* multiple comparisons showed that CaMKII-mCherry mice increased their average freezing levels between the 3 vs. 9 month (Tukey’s: p=0.0008) and 3 vs. 6 month (Tukey’s: p=0.0089) time points, indicating an aging effect (Figure 2C; iv). Most interestingly, at the 3 month time point there is a significant increase in average freezing in the CaMKII-hM3Dq mice compared to the mCherry controls, indicating that our manipulation is affecting extinction behavior at this early time point. Within session at 3 months, there was no interaction between time bin and group, but there were significant differences in freezing depending on time bin and group alone (Figure 2C; i)(Two-way ANOVA RM; Interaction: F (29, 580) = 0.5858, p=0.9601; Time: F (7.717, 154.3) = 2.642, p=0.0105; Group: F (1, 20) = 8.560, p=0.0084; Subject: F (20, 580) = 12.70, p<0.0001). At 6 months, there was no significant interaction between time bin and group, but there was an effect of time only on freezing levels during extinction day 1 (Figure 2C; ii)(Two-way ANOVA RM; Interaction: F (29, 638) = 0.4496, p=0.9948; Time: F (8.852, 194.7) = 6.720, p<0.0001; Group: F (1, 22) = 0.3101, p=0.5832; Subject: F (22, 638) = 19.17, p<0.0001). Finally, at 9 months there was no interaction between time bin and group, nor was freezing impacted by time bin or group alone (Figure 2C; iv)(Mixed-effects model (REML); Time: F (6.259, 110.1) = 2.097, p=0.0565; Group: F (1, 18) = 3.598, p=0.0740; Interaction: F (29, 510) = 1.342, p=0.1121)(Figure 2C; iii).

For extinction day 2, there was a significant interaction between time point and group for average freezing (Two-way ANOVA: Interaction: F(2,58) = 4.511, p=0.0151; Time: F(2,58) = 2.760, p=0.0716; Group: F(1,58) = 0.3378, p=0.5633)(Figure 2D; iv). *Post hoc* analysis revealed that CaMKII-mCherry mice have an increase in freezing level across the 3 and 6 month time points with aging (Tukey’s: p=0.0464). Within the 3 month time point, there was no interaction between time bin and group, but there were effects of time and group alone on the percent freezing within session (Two-way ANOVA RM; Interaction: F (29, 580) = 1.298, p=0.1385; Time: F (6.958, 139.2) = 3.593, p=0.0014; Group: F (1, 20)= 6.345, p=0.0204; Subject: F (20, 580) = 15.50, p<0.0001) (Figure 2D; i). Within session at 6 months, there was no interaction within session between time bin and group, however, time bin did have an effect on the difference in freezing during extinction day 2 (Two-way ANOVA RM; Interaction: F (29, 696) = 1.130, p=0.2921; Time: F (5.396, 129.5) = 2.888, p=0.0142; Group: F (1, 24) = 1.313, p=0.2632; Subject: F (24, 696) = 35.67, p<0.0001) (Figure 2D; ii). At 9 months there was a significant interaction between time bin and group on freezing levels within session for extinction day 2 (Two-way ANOVA RM; Interaction: F (29, 522) = 1.918, p=0.0031; Time: F (6.146, 110.6) = 2.383, p=0.0322; Group: F (1, 18) = 0.1419, p=0.7108; Subject: F (18, 522) = 16.26, p<0.0001)(Figure 2D; iii). *Post-hoc* multiple comparisons did not reveal pairwise differences across groups for each time bin.

For extinction day 3, there was no interaction between time point and group, but there was an effect of time point on average freezing levels (Two-way ANOVA; Interaction: F (2, 60) = 2.147, p=0.1257; Timepoint: F (2, 60) = 7.099, p=0.0017; Group: F (1, 60) = 0.006334, p=0.9368) (Figure 2E; iv). Within session at 3 months, there was no significant interaction between time bin and group (Two-way ANOVA RM; Interaction: F(29,580) = 0.5950, p=0.9557; Time: F(4.761,95.23)=1.417, p=0.2273; Group: F(1,20) = 3.648, p=0.0706; Mouse: F(20,580) = 14.34, p<0.0001)(Figure 2E; i). Within session at 6 months, there was a significant interaction between time bin and group, but *post hoc* multiple comparisons did not reveal pairwise differences (Two-way ANOVA RM; Interaction: F (29, 638) = 1.583, p=0.0277; Time: F (6.744, 148.4) = 3.160, p=0.0043; Group: F (1, 22) = 2.086, p=0.1627; Subject: F (22, 638) = 27.14, p<0.0001)(Figure 2E; ii). Finally, within sessions at the 9 month time point there was no significant interaction between time bin and group (Two-way ANOVA RM; Interaction: F(29,522) = 0.7420, p=0.8353; Time: F(7.702, 138.6) = 1.332, p=0.2345; Group: F(1,18) = 0.06444, p=0.8025; Mouse: F(18,522) = 19.96, p<0.0001)(Figure 2E; iii).

When observing behavior across days, CaMKII-hM3Dq and mCherry groups had no interaction between day [fear conditioning, recall, extinction days 1-3] and group [CaMKII-hM3Dq and mCherry] at the 3, 6 and 9 month time points. However, at the 3 and 9 month time points, there were individual effects of both day and group, suggesting that our manipulation of the Gq pathway was impacting average freezing levels. Additionally, at the 6 month time point, there was an effect of day only, suggesting that our experimental manipulation was not having an impact at this time point (Two-way ANOVA RM; [3 Month: Interaction: F (4, 80) = 1.166, p=0.3320; Day: F (2.954, 59.08) = 42.66, p<0.0001; Group: F (1, 20) = 5.443, p=0.0302; Subject: F (20, 80) = 5.297, p<0.0001][6 month: Interaction: F (4, 88) = 0.5515, p=0.6984; Day: F (4, 88) = 25.72, p<0.0001; Group: F (1, 22) = 2.571, p=0.1231; Subject: F (22, 88) = 1.949, p=0.0153][9 month: Interaction: F (4, 72) = 1.580, p=0.1888; Day: F (2.640, 47.52) = 28.45, p<0.0001; Group: F (1, 18) = 11.43, p=0.0033; Subject: F (18, 72) = 1.158, p=0.3188])(Figure 2F-H).

In summary, our CaMKII-hM3Dq manipulation significantly decreased freezing levels during CFC compared to CaMKII-mCherry controls at the 9 month time point. During CFC, we also observed an increase in fear with aging in the 3 vs. 9 month CaMKII-mCherry groups. For recall, we observed a change in average freezing due to aging alone. For extinction day 1, we observed effects of aging across the 3 vs. 9 month and 3 vs. 6 month CaMKII-mCherry groups. Most notably, we observed a significant increase in average freezing during extinction in CaMKII-hM3Dq compared to control mice at the 3 month time point. For extinction day 2, there were individual impacts of aging and our manipulation for average freezing levels. Specifically, we observed an increase in freezing across 3 vs. 6 month CaMKII-mCherry mice. Finally, for extinction day 3 we only observed an impact of aging on freezing behavior. Our findings that Gq activation increases freezing at the 3 month time point during extinction may indicate that our intervention increases anxiety and/or impairs extinction mechanisms. Additionally, a decrease in freezing at the 9 month time point during CFC may indicate a decrease in anxiety or impairments in memory encoding. Overall, our neuronal-targeted Gq manipulation produces differential effects on contextual fear acquisition, recall and extinction behaviors, suggesting that normal functioning of excitatory neurons in vHPC is necessary for proper fear acquisition and maintenance.

### 3.3) Chronic Gq activation of GFAP+ astrocytes mildly impacts freezing behavior during contextual fear acquisition and maintenance

Next, to test the hypothesis that chronic astrocytic Gq activation is sufficient to induce behavioral deficits across all timepoints, we subjected our mice to CFC, recall, and three days of extinction (Figure 1E). For CFC, there was a significant interaction between group and time point for average freezing, suggesting that our manipulation and aging are interacting to impact memory acquisition (Two-way ANOVA; Interaction: F(2,56) = 3.24, p=0.0466; Timepoint: F (2, 56) = 0.6979, p=0.05019; Group: F (1, 56) = 0.8864, p=0.3505)(Figure 3A; iv). *Post hoc* multiple comparisons did not reveal significant pairwise differences between groups (Figure 3A; iv). Within the session, mice at 3 months showed no interaction between time bin and group. On the other hand, each factor alone had an effect on freezing level during CFC (Two-way ANOVA RM; Interaction: F (5, 115) = 1.201, p=0.3133; Time: F (5, 115) = 135.1, p<0.0001; Group: F (1, 23) = 8.509, p=0.0078; Mouse: F (23, 115) = 2.421, p=0.0011)(Figure 3A; i). At 6 months, there was no interaction between time bin and group, but there was an effect of time bin alone that is expected as mice acquire fear successfully within session (Two-way ANOVA RM; Interaction: F (5, 85) = 1.054, p=0.3916; Time: F (3.198, 54.36) = 71.62, p<0.0001; Group: F (1, 17) = 0.1673, p=0.6877; Mouse: F (17, 85) = 3.188, p=0.0002)(Figure 3A; ii). At 9 months, there was no significant interaction between time bin and group within session (Two-way ANOVA RM; Interaction: F (5, 90) = 0.2597; Time: F (2.988, 53.79) = 36.41, p<0.0001; Group: F (1, 18) = 3.339, p=0.0843; Mouse: F (18, 90) = 3.201, p=0.0001)(Figure 3A; iii).

**Figure 3.**
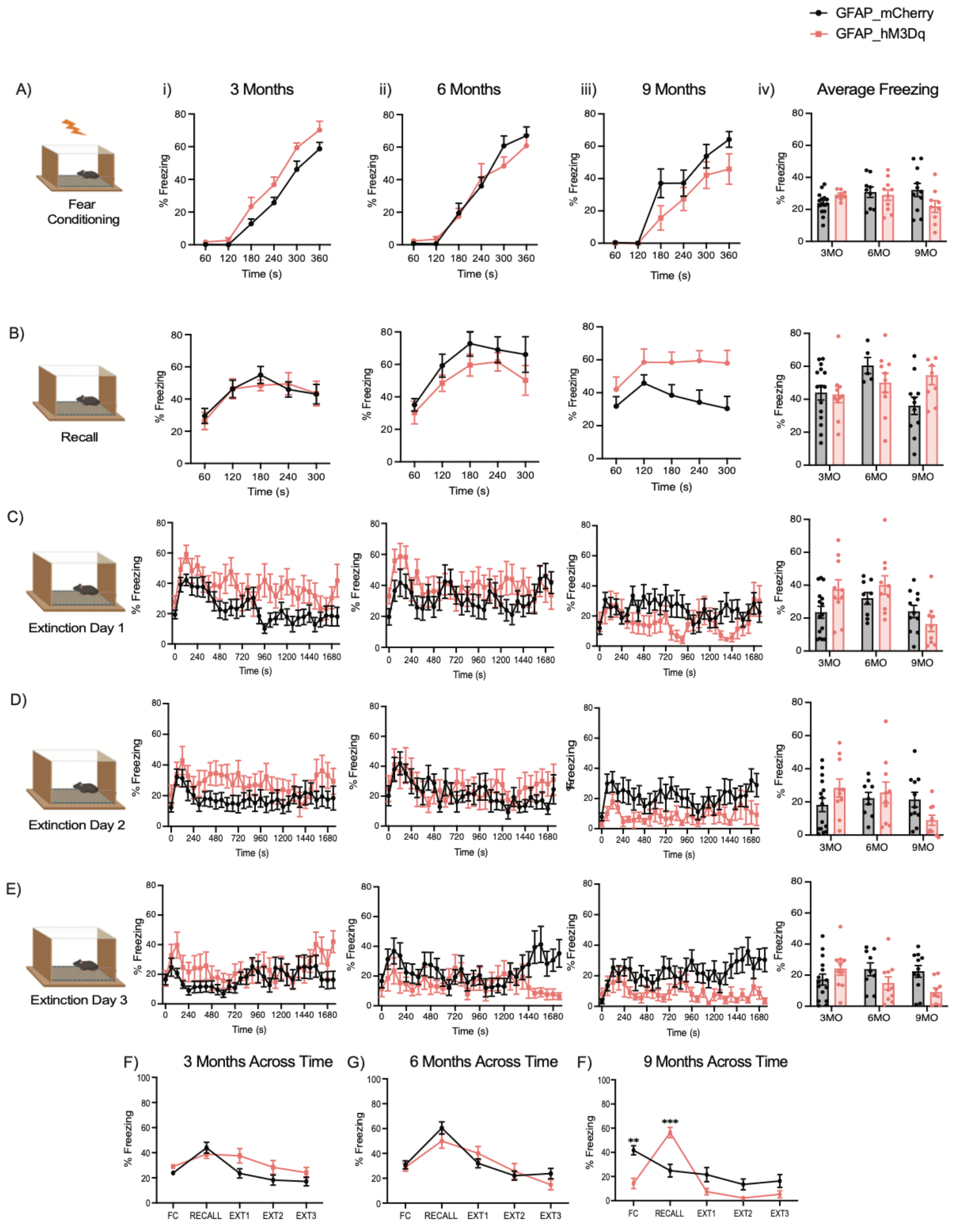
Gq pathway activation in GFAP+ astrocytes mildly impacts CFC, recall and extinction at the 6 and 9 month time points. (A-E) Percent freezing levels and freezing percentage across time (seconds) of 3, 6, and 9 month time points. (A) GFAP groups during contextual fear conditioning (CFC), where mice received four, 1.5mA foot shocks at 120, 180, 240 and 300 seconds during a 360 second session, (B) recall session for 360 seconds, where mice did not receive the aversive stimulus, (C) extinction day 1, (D) day 2, and (E) day 3, where mice received no foot shock for a 1800 second session. (F-H) Average percent freezing (%) across fear conditioning, recall and extinction days 1-3 for GFAP-hM3Dq and mCherry groups at 3 month (F), 6 month (G) and 9 month (H) time points. Average percent freezing was assessed with a two-way analysis of variance (ANOVA) with time point and group as factors. Freezing percentage across the 1-minute time bins [60, 120, 180, 240, 300, 360] was assessed with a two-way ANOVA with repeated measures (RM) with time (seconds) and group as factors. Average percent freezing across days [FC, Recall, EXT1, EXT2, EXT3] was assessed with a two-way ANOVA with repeated measures (RM) with day and group as factors. Tukey’s and Sidak’s post hoc tests were performed where applicable. Error bars indicate SEM. p ≤ 0.05, **p ≤ 0.01, ***p ≤ 0.001, ****p ≤ 0.0001, ns = not significant. 3 Month: hM3Dq (n=8-10), mCherry (n=15); 6 Month: hM3Dq (n=10), mCherry (n=9); 9 Months: hM3Dq (n=9), mCherry (n=11) after outlier removal.

For recall, there was a significant interaction between time point and group (Two-way ANOVA; Interaction: F (2, 54) = 3.357, p=0.0422; Timepoint: F (2, 54) = 2.267, p=0.1134; Group: F (1, 54) = 0.2532, p=0.6169)(Figure 3B; iv). *Post hoc* analysis did not reveal significant differences between groups or across time points. Within session at 3 and 6 months, there was no interaction between time bin and group, but there was an effect of time bin alone (Two-way ANOVA RM; [3 Month: Interaction: F (4, 92) = 0.5448, p=0.7033; Time: F (4, 92) = 12.54, p <0.000; Group: F (1, 23) = 0.02859, p=0.867; Mouse: F (23, 92) = 8.422, p<0.0001)][6 Month: Interaction: F (4, 52) = 0.3647, p=0.8326; Time: F (2.512, 32.65) = 13.56, p<0.0001; Group: F (1, 13) = 1.326, p=0.2702; Mouse: F (13, 52) = 7.540, p<0.0001])(Figure 3B; i, ii). Finally, at 9 months there was no interaction between time bin and group within session, but group alone did have an effect on freezing levels within the recall session (Two-way ANOVA RM; Interaction: F (4, 72) = 1.389; p=0.2464; Time: F (2.131, 38.35) = 3.132, 0.0520; Group: F (1, 18) = 6.032, p=0.0244; Mouse: F(18,72) =7.314, p<0.0001)(Figure 3B; iii).

We observed no significant interaction between time point and group for extinction day 1, only an effect of time point alone, suggesting that only aging contributed to the variance in freezing behavior (Two-way ANOVA; Interaction: F (2, 58) = 3.041, p=0.0555; Timepoint: F (2, 58) = 5.841, p=0.0049; Group: F (1, 58) = 1.637, p=0.2059)(Figure 3C; iv). Within session at 3 months, there was no interaction between time bin and group, however each factor contributed independently to freezing levels during extinction (Two-way ANOVA RM; Interaction: F (29, 690) = 0.6285, p=0.9370; Time: F (29, 690) = 2.529, p<0.0001; Group: F (1, 690) = 63.51, p<0.0001)(Figure 3C; i). At 6 and 9 months, there was no interaction between time bin and group, but our manipulation alone contributed significantly to freezing levels within session (Two-way ANOVA; [6 Months: Interaction: F (29, 510) = 0.5914, p=0.9571; Time: F (29, 510) = 0.9590, p=0.5292; Group: F (1, 510) = 14.36, p=0.0002][9 Months: Interaction: F (29, 536) = 0.9447, p=0.5508; Time: F (29, 536) = 0.8777, p=0.6527; Group: F (1, 536) = 20.73, p<0.0001])(Figure 3C; ii, iii).

For extinction day 2, there was no interaction between group and time point for average freezing levels (Two-way ANOVA; Interaction: F (2, 58) = 3.073, p=0.0539; Time point: F (2, 58) = 2.062, p=0.1365; Group: F (1, 58) = 0.01206, p=0.9129)(Figure 3D; iv). Within session at 3 months, there was no interaction between time bin and group, but time bin alone had an effect on freezing across extinction (Two-way ANOVA RM; Interaction: F (29, 667) = 0.9633, p=0.5225; Time: F (29, 667) = 1.735, p=0.0102; Group: F (1, 23) = 2.356, p=0.1385; Mouse: F (23, 667) = 29.95, p<0.0001)(Figure 3D; i). At 6 and 9 months, there was no significant interaction between time bin and group, only an effect of group at the 9 month time point (Two-way ANOVA RM; [6 Month: Interaction: F (29, 493) = 0.9984, p=0.4704; Time: F (6.484, 110.2) = 2.020, p=0.0637; Group: F (1, 17) = 0.2175, p=0.6468; Mouse: F (17, 493) = 23.34, p<0.0001)][9 Month: Interaction: F (29, 522) = 0.5766, p=0.9640; Time: F (5.557, 100.0) = 1.231, p=0.2986; Group: F (1, 18) = 4.786, p=0.0421; Mouse: F (18, 522) = 18.83, p<0.0001])(Figure 3D, ii-iii).

For extinction day 3 there was an interaction between group and time point for average freezing levels, suggesting that age and our manipulation impacted behavior (Two-way ANOVA; Interaction: F (2, 58) = 4.106, p=0.0215; Timepoint: F (2, 58) = 0.8808, p=0.4199; Group: F (1, 58) = 2.624, p=0.1107)(Figure 3E, iv). *Post hoc* multiple comparisons did not reveal significant differences in means across groups or time points. At 3 months, there was no significant interaction between time bin and group within session, but time bin had an effect on the change in freezing level (Two-way ANOVA RM; Interaction: F (29, 667) = 1.078, p=0.3573; Time: F (6.706, 154.2) = 2.136, p=0.0455; Group: F (1, 23) = 1.622, p=0.2155; Mouse: F (23, 667) = 16.12, p<0.0001)(Figure 3E; i). However, at 6 months there was an interaction between group and time bin on freezing levels within session, suggesting that our manipulation had an effect in the GFAP-hM3Dq group (Two-way ANOVA RM; Interaction: F (29, 493) = 1.857, p=0.0048; Time: F (7.353, 125.0) = 1.200, p=0.3067; Group: F (1, 17) = 2.354, p=0.1433; Mouse: F (17, 493) = 16.51, p<0.0001)(Figure 3E; ii). Finally, at 9 months there was no significant interaction between time bin and group, but there was a main effect of group on freezing levels within session (Two-way ANOVA RM; Interaction: F (29, 522) = 1.423, p=0.0726; Time: F (6.763, 121.7) = 1.057, p=0.3948; Group: F (1, 18) = 7.594, p=0.0130; Mouse: F (18, 522) = 14.46, p=0.0130)(Figure 3E, iii).

When measuring across days, GFAP-hM3Dq and mCherry groups had no significant interaction between day and group at the 3 and 6 month time points. However, there was an effect of day alone, suggesting that our manipulation was not having any effect on learning (Mixed-effects model (REML): [3 Month: Interaction: F (4, 89) = 2.096, p=0.0879; Day; F (4, 89) = 11.18, p<0.0001; Group: F (1, 23) = 2.859, p=0.1044][6 Month: Interaction: F (4, 81) = 1.342, p=0.2617; Day: F (4, 81) = 14.72, p<0.0001; Group: F (1, 81) = 0.4120, p=0.5228])(Figure 3F-G). Most interestingly, there was a significant interaction between day and group at the 9 month time point for GFAP-hM3Dq and mCherry groups (Two-way ANOVA RM; Interaction: F (4, 72) = 6.130, p=0.0003; Day: F (4, 72) = 21.41, p<0.0001; Group: F (1, 18) = 1.948, p=0.1798; Mouse: F (18, 72) = 2.259, p=0.0079)(Figure 3H). Further *post hoc*analysis revealed significant differences in freezing between groups at the 9 month time point for the recall day (Sidak’s: p=0.0142).

In summary, we observed significant individual impacts of our manipulation and aging on average freezing during CFC. Only the 3 month time point showed an impact of our manipulation on within-session freezing levels during CFC. For recall, there was a significant impact of both aging and our manipulation on average freezing levels. Only the 9 month time point showed an impact of our manipulation on within-session freezing levels during contextual recall. For extinction day 1, there was only an effect of aging on average freezing levels. All of the time points showed an impact of our manipulation on within-session freezing levels during extinction day 1. For extinction day 2, there was no impact of aging or our manipulation on average freezing levels. Only the 9 month time point demonstrated an effect of the group on freezing levels within-session for extinction day 2. Finally, for extinction day 3 there was an impact of both group and aging on average freezing levels. At 6 and 9 months within-session, there was a difference in freezing levels that could be described by group manipulation.

### 3.4) Chronic Gq activation of neurons induces behavioral changes in locomotor and anxiety-related behaviors

The open field and elevated zero maze are tasks that are classically used to measure locomotor activity and anxiety-like behaviors (Seibenhener & Wooten, 2015; Tucker & McCabe, 2017). To test our hypothesis that chronic activation of the Gq pathway in neurons will exhibit disrupted anxiety-like behaviors, we subjected the mice to these tasks on consecutive days (Figure 1E). For open field, CaMKII-hM3Dq and mCherry mice had no significant interaction between time point and group in the total distance traveled, time spent in center, and number of entries to the center (Figure 4A, C-D). However, each factor alone (i.e. group and time point) individually contributed to all three of these metrics (Two-way ANOVA; [Distance traveled: Interaction: F (2, 65) = 0.5961, p=0.5540; Timepoint: F (2, 65) = 3.437, p=0.0381; Group: F (1, 65) = 6.664, p=0.0121][Center time: Interaction: F (2, 56) = 0.5527, p=0.5785; Timepoint: F (2, 56) = 13.97, p<0.0001; Group: F (1, 56) = 7.767, p=0.0073][Center entries: Interaction: F (2, 58) = 1.353, p=0.2664; Timepoint: F (2, 58) = 8.707, p=0.0005; Group: F (1, 58) = 4.134, p=0.0466])(Figure 4A, C-D). This suggests that both aging and our Gq pathway manipulation are individually driving differences in anxiety-related behaviors. Mean speed in the open field demonstrated no interaction between time point and group, but group alone contributed to a modification in behavior at each time point (Two-way ANOVA; Interaction: F (2, 62) = 0.2549, p=0.7758; Time point: F (2, 62) = 3.103, p=0.0519; Group: F (1, 62) = 4.937, p=0.0299)(Figure 4B). Further, there was no significant interaction between time point and group for center mean visit time (Two-way ANOVA; Interaction: F (2, 57) = 1.168, p=0.3182; Timepoint: F (2, 57) = 1.972, p=0.1486; Group: F (1, 57) = 2.985, p=0.08950)(Figure 4E). This suggests that the combination of aging and the manipulation of the CaMKII-Gq pathway drives significant differences in anxiety-related behaviors.

**Figure 4.**
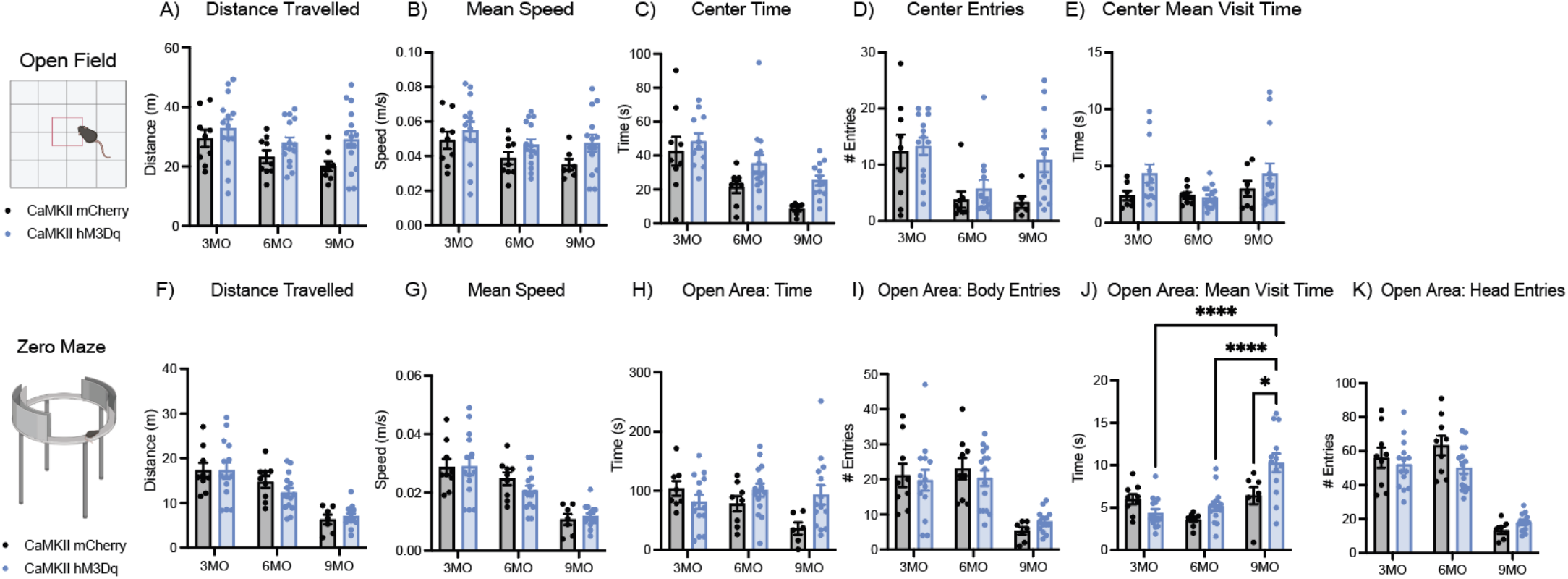
Chronic Gq activation of neurons and aging induces behavioral changes in locomotor and anxiety-related behaviors. (A-E) Open field distance traveled (A), mean speed (B), total time spent in center (C), number of entries into center (D), and mean center visit time (E) in 3, 6, and 9 month CaMKII groups. (F-M) Zero maze distance traveled (F), mean speed (G), total time spent in open area (H), number body entries into open area (I), mean open area visit time (J), and number of head entries into open area (K) in 3, 6, and 9 month CaMKII groups. Open field and zero maze metrics were assessed with a two-way analysis of variance (ANOVA) with time point and group as factors. Tukey’s post hoc tests were performed where applicable. Error bars indicate SEM. p ≤ 0.05, **p ≤ 0.01, ***p ≤ 0.001, ****p ≤ 0.0001, ns = not significant. For open field, 3 Month: hM3Dq (n=11-14), mCherry (n=7-9); 6 Months: hM3Dq (n=13-15), mCherry (n=8-9); 9 Months: hM3Dq (n=12-15), mCherry (n=6-9) after outlier removal. For zero maze, 3 Month: hM3Dq (n=14), mCherry (n=9); 6 Month: hM3Dq (n=15), mCherry (n=8-9); 9 Month: hM3Dq (n=14), mCherry (n=6-7) after outlier removal.

In the elevated zero maze, there was no significant interaction between time point and group for total distance traveled, mean speed, body entries into the open area, or head entries into the open area. However, time point alone had an effect on behavior for all of these metrics, with each measure differing across aging (Two-way ANOVA; [Distance traveled: Interaction: F (2, 62) = 0.6878, p=0.5065; Timepoint: F (2, 62) = 27.98, p<0.0001; Group: F (1, 62) = 0.2502, p=0.6187][Mean speed: Interaction: F (2, 62) = 0.7047, p=0.4982; Timepoint: F (2, 62) = 27.25, p<0.0001; Group: F (1, 62) = 0.2245, p=0.6373][Open area body entries: Interaction: F (2, 62) = 0.5662, p=0.5706; Timepoint: F (2, 62) = 19.92, p<0.0001; Group: F (1, 62) = 0.04116, p=0.8399][Open area head entries: Interaction: F (2, 62) = 2.369, p=0.1020; Timepoint: F (2, 62) = 60.87, p<0.0001; Group: F (1, 62) = 1.614, p=0.2086])(Figure 4F-G, I, K). Interestingly, for the time spent in the open area and mean visit time there was an interaction between both time point and our group manipulation (Two-way ANOVA; [Open area time: Interaction: F (2, 59) = 3.564, p=0.0346; Timepoint: F (2, 59) = 2.028, p=0.1406; Group: F (1, 59) = 2.514, p=0.1182][Open area mean visit time: Interaction: F (2, 61) = 6.537, p=0.0027; Timepoint: F (2, 61) = 14.68, p<0.0001; Group: F (1, 61) = 4.244, p=0.0437])(Figure 4H, J). *Post hoc* multiple comparisons revealed significant pairwise increases across 3 vs. 9 month CaMKII-hM3Dq (Tukey’s: p<0.0001) and the 6 vs. 9 month CaMKII-hM3Dq groups (Tukey’s: p<0.0001), with open area mean visit time increasing with age (Figure 4J). Most interestingly, our manipulation of the CaMKII-Gq pathway induced a significant increase in the mean visit time to the open area of the zero maze at the 9 month time point when comparing CaMKII-hM3Dq and mCherry groups (Tukey’s: p=0.0148)(Figure 4J). Finally, we observe normal aging changes in distance traveled, mean speed, body entries and head entries to the open area, indicating an increase in anxiety with age (Yanai & Endo, 2021). Thus overall, CaMKII Gq activation had a combinatory effect on anxiety-related behaviors with age.

### 3.5) Chronic Gq activation of astrocytes induces changes in locomotion and anxiety-related behaviors, as evidenced by the open field and zero maze tests

We next tested the hypothesis that chronic modulation of the Gq pathway in astrocytes would induce changes in anxiety-related behaviors (Figure 1E). For open field, GFAP-hM3Dq and mCherry mice showed no significant interaction between time point and group in the total distance traveled and mean speed. However, there was an effect of time point alone for both measures, indicating an effect on distance traveled and mean speed with aging that is shown in previous literature (Two-way ANOVA; [Distance Traveled: Interaction: F (2, 57)= 0.3488, p=0.7070; Timepoint: F (2, 57) = 14.98, p<0.0001; Group: F (1, 57) = 0.006246, p-0.9373][Mean speed: Interaction: F (2, 57) = 0.3533, p=0.7039; Timepoint: F (2, 57) = 14.97, p<0.0001; Group: F (1, 57) =0.003799, p=0.9511])(Figure 5A-B)(Yanai & Endo, 2021). For time spent in the open field, there was a significant interaction between time point and group for GFAP-hM3Dq and mCherry mice (Two-way ANOVA; Interaction: F (2, 55) = 4.012, p=0.0236; Timepoint: F(2,55) = 5.135, p=0.0090; Group: F(1,55) = 2.519, p=0.1182)(Figure 5C). *Post hoc* multiple comparisons revealed a significant increase in time spent in the center for GFAP-hM3Dq mice at the 3 month time point (Tukey’s: p=0.0192), indicating a decrease in anxiety-like behavior due to our manipulation (Figure 5C). Additionally, 3 vs. 6 month GFAP-hM3Dq mice exhibited a decrease in the time spent in the center of the open field (Tukey’s: p=0.0305), suggesting that aging in combination with our manipulation is having an effect on anxiety-related behaviors (Figure 5C). For the number of entries into the center of the open field, there was no interaction between time point and group, only an individual effect of time point alone, suggesting that aging is having an impact on anxiety-related behaviors (Two-way ANOVA; Interaction: F (2, 63) = 2.586, p=0.0833; Timepoint: F (2, 63) = 3.719, p=0.0297; Group: F (1, 63) = 0.1661, p=0.6850)(Figure 5D). Finally, for the mean visit time to the center of the open field, there was no interaction between time point and group, but time point and group individually contributed to changes in this measure of anxiety related behavior (Two-way ANOVA; Interaction: F (2, 48) = 0.7735, p=0.4670; Timepoint: F (2, 48) =3.773, p=0.0301; Group: F (1, 48) =8.772, p=0.0049)(Figure 5E). In summary, there was an expected decrease in distance traveled and mean speed with aging, consistent with previous literature (Yanai & Endo, 2021; Shoji & Miyakawa, 2019). Evidence of aging effects were also present in the mean visit time to the center of the open field, with time point having an effect on the typical amount of time the mouse spent in the open area. Our manipulation of the Gq pathway in GFAP+ cells induced notable increases in the time spent in the center of the open field at the 3 month time point, indicating that vHPC manipulation disrupts anxiety-related behaviors. Additionally, in time spent in the center of the open field, there was a decrease in time between 3 and 6 months of age for the GFAP-hM3Dq group, indicating an effect of aging that is likely being amplified by our manipulation.

**Figure 5.**
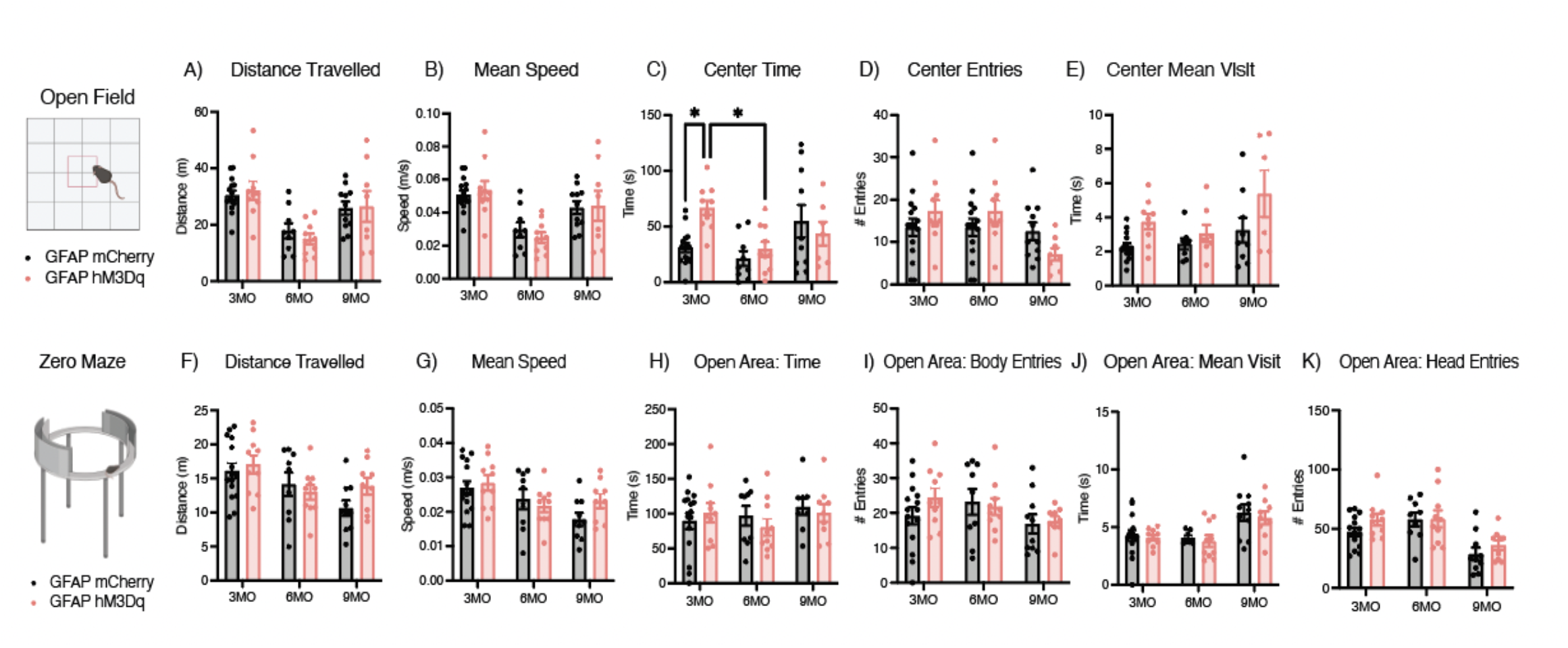
Chronic Gq activation of astrocytes decreases locomotor and anxiety-related behaviors in the open field at the 3 month time point, but only aging affects zero maze behavior. (A-E) Open field distance traveled (A), mean speed (B), total time spent in center (C), number of entries into center (D), and mean center visit time (E) in 3, 6, and 9 month GFAP groups. (F-M) Zero maze distance traveled (F), mean speed (G), total time spent in open area (H), number body entries into open area (I), mean open area visit time (J), and number of head entries into open area (K) in 3, 6, and 9 month GFAP groups. Open field and zero maze metrics were assessed with a two-way analysis of variance (ANOVA) with time point and group as factors. Tukey’s post hoc tests were performed where applicable. Error bars indicate SEM. p ≤ 0.05, **p ≤ 0.01, ***p ≤ 0.001, ****p ≤ 0.0001, ns = not significant. For open field, 3 Month: hM3Dq (n=9-10), mCherry (n=14-15); 6 Month: hM3Dq (n=8-10), mCherry (n=8-9); 9 Month: hM3Dq (n=6-8), mCherry (n=9-11) after outlier removal. For zero maze, 3 Month: hM3Dq (n=10), mCherry (n=15); 6 Month: hM3Dq (n=10), mCherry (n=9); 9 Month: hM3Dq (n=9), mCherry (n=10) after outlier removal.

For elevated zero maze, GFAP-hM3Dq and mCherry mice showed no significant interaction between time point and group in the total distance traveled, mean speed, number of body entries, number of head entries, time spent and mean visit time to the open area of the apparatus. However, distance traveled, mean speed, head entries and mean visit time displayed an individual effect of time point, suggesting that aging is impacting some aspects of anxiety-related behaviors in this task (Two-way ANOVA; [Distance traveled: Interaction: F (2, 57) = 1.334, p=0.2716; Timepoint: F (2, 57) = 6.309, p=0.0033; Group: F (1, 57) = 0.8936, p=0.3485][Mean speed: Interaction: F (2, 57) = 1.313, p=0.2770; Timepoint: F (2, 57) = 6.203, p=0.0037; Group: F (1, 57) = 0.8219, p=0.3684][Open area body entities: Interaction: F (2, 57) = 0.8330, p=0.4400; Timepoint: F (2, 57) = 2.103, p=0.1314; Group: F (1, 57) = 0.4727, p=0.4945][Open area head entries: Interaction: F (2, 57) = 0.4850, p=0.6182; Timepoint: F (2, 57) = 12.76, p<0.0001; Group: F (1, 57) = 2.049, p=0.1578][Open area time: Interaction: F (2, 56) = 0.6650, p=0.5183; Timepoint: F (2, 56) = 0.7516, p=0.4763; Group: F (1, 56) = 0.1791, p=0.6737][Open area mean visit time: Interaction: F (2, 57) = 0.04414, p=0.9569; Timepoint: F (2, 57) = 9.812, p=0.0002; Group: F (1, 57) = 0.5125, p=0.4770])(Figure 5F-K). Overall, our manipulation of the Gq pathway in astrocytes impacts anxiety-related behaviors in the open field, but not in the elevated zero maze. Most notably, we observe a decrease in anxiety at the 3 month time point, as evidenced by an increase in the time spent in the center of the open field.

### 3.6) Chronic Gq activation impacts social behaviors in CaMKII- and GFAP-hM3Dq groups

To assess how social behaviors are impacted by our manipulation, we performed a social interaction test by placing the subject mouse in an open arena with a cup containing a male mouse or an empty cup for a 10 minute session. For CaMKII-hM3Dq and mCherry mice, there was no interaction between time point and group for total distance traveled, head time spent at the empty cup, or the number of head entries into the empty cup area. However, these metrics were all impacted by an effect of time point alone, suggesting that aging is contributing to these behavioral changes (Two-way ANOVA; [Distance traveled: Interaction: F (2, 59) = 1.447, p=0.2435; Timepoint: F (2, 59) = 69.58, p<0.0001; Group: F (1, 59) = 0.1835, p=0.6699][Empty cup head time: Interaction: F (2, 59) = 2.514, p=0.0896; Time point: F (2, 59) = 13.96, p<0.0001; Group: F (1, 59) = 0.0005010, p=0.9822][Empty cup mean visit time: Interaction: F (2, 55) = 0.8991, p=0.4128; Timepoint: F (2, 55) = 30.29, p<0.0001; Group: F (1, 55) = 0.7430, p=0.3924])(Supplemental Figure 1D-F). Interestingly, for the number of head entries into the mouse cup (i.e. social target), there was a significant interaction between time point and group (Two-way ANOVA; Interaction: F (2, 59) = 6.145, p=0.0038; Timepoint: F (2, 59) = 35.11, p<0.0001; Group: F (1, 59) = 0.4773, p=0.4924)(Supplemental Figure 1A). *Post hoc* analysis revealed significant decreases in the number of head entries into the mouse cup between 3 vs. 9 month CaMKII-mCherry (Tukey’s: p<0.0001), 6 vs. 9 month CaMKII-mCherry (Tukey’s: p<0.0001), 3 vs. 9 month CaMKII-hM3Dq (Tukey’s: p=0.0002), and 6 vs. 9 month CaMKII-hM3Dq (Tukey’s: p=0.0176) groups (Supplemental Figure 1A). Most notably, there was a significant decrease in the number of head entries into the mouse cup at the 6 month time point between our CaMKII-hM3Dq and mCherry groups (Tukey’s: p=0.0391)(Supplemental Figure 1A). Finally, for the total amount of head time and mean visit time to the mouse cup, as well as the mean visit time to the empty cup, there was no significant interaction between time point and group (Two-way ANOVA; [Mouse cup head time: Interaction: F (2, 56) = 1.056, p=0.3547; Time point: F (2, 56) = 0.5529, p=0.5784; Group: F (1, 56) = 0.8246, p=0.3677][Mouse cup mean visit time: Interaction: F (2, 55) = 1.138, p=0.3278; Timepoint: F (2, 55) = 2.494, p=0.0919; Group: F (1, 55) = 3.475, p=0.0676][Empty cup mean visit time: Interaction: F (2, 52) = 1.333, p=0.2727; Timepoint: F (2, 52) = 2.111, p=0.1314; Group: F (1, 52) = 0.7496, p=0.3906])(Supplemental Figure 1B-C, G).

Overall, we found a significant decrease in the number of entries into the mouse cup area within the 6 month time point, suggesting that these mice may have decreased interest in the social target with our manipulation. CaMKII-hM3Dq and mCherry mice displayed no significant differences in distance traveled, time spent at the empty cup and head entries into the empty cup zone, but aging did have an impact. Head entries into the mouse cup had a significant impact of aging and our manipulation, with a decrease in the number of entries across the 3 vs. 9 month and 6 vs. 9 month time points for both groups. These results are consistent with effects of aging, such as general decreased interest in social targets (Shoji et al, 2016; Oizumi et al, 2019; Yanai & Endo, 2021).

For GFAP-hM3Dq and mCherry mice, there was no significant interaction between time point and group for total distance traveled, mean speed, head time at the mouse cup (i.e. social target) or empty cup (i.e. control), head entries into the mouse cup or empty cup zones, or mean visit time to the mouse cup or empty cup during social interaction. However, we did observe significant effects of aging across all mouse cup metrics, but not in the empty cup metrics (Two-way ANOVA; [Distance traveled: Interaction: F (2, 56) = 1.617, p=0.2077; Timepoint: F(2,56) = 7.854, p=0.0010; Group: F(1,56) = 1.219, p=0.2743][Mean speed: Interaction: F(2,55) = 1.485, p=0.2355; Timepoint: F(2,55) = 7.413, p=0.0014; Group: F(1,55) = 1.291, p=0.2609][Mouse cup head time: Interaction: F(2,52) = 1.792, p=0.1767; Timepoint: F(2,52) = 52.18, p<0.0001; Group: F(1,52) = 0.5226, p=0.4730][Mouse cup head entries: Interaction: F (2, 55) = 0.4886, p=0.6161; Timepoint: F(2,55) = 15.79, p<0.0001; Group: F(1,55) = 0.7936, p=0.3769][Mouse cup mean visit time: Interaction: F(2,55) = 1.336, p=0.2714; Timepoint: F(2,55) = 3.480, p=0.0377; Group: F(1,55) = 1.012, p=0.3189][Empty cup head time: Interaction: F(2,56) = 1.028, p=0.3644; Timepoint: F(2,56) = 2.460, p=0.0947; Group: F(1,56) = 0.4234, p=0.5179][Empty cup head entries: Interaction: F(2,56) = 0.4035, p=0.6699; Timepoint: F(2,56) = 2.542, p=0.0878; Group: F(1,56) = 0.007, p=0.9329][Empty cup mean visit time: Interaction: F(2,55) = 0.3364, p=0.7158; Timepoint: F(2,55) = 1.977, p=0.1482; Group: F(1,55) = 0.3792, p=0.5406](Supplemental Figure 2A-G). Overall, we observed effects of aging on measures of locomotion (i.e. distance traveled and mean speed) and engagement levels with the social target (i.e. mouse cup), but not in the control target (i.e. empty cup) metrics.

### 3.7) Novel environment exploration is affected by CaMKII- and GFAP-hM3Dq activation

When exploring a novel environment, rodents generally investigate novel objects and spaces. To assess novel context exploration and quantify putative cognitive deficits, we performed an elevated y-maze test. Our assessment of novel environment exploration using the y-maze can also be used to assess short-term spatial working memory, as mice are required to remember the most recent arm they explored and navigate to a ‘novel’ arm instead. After introduction into the center of the y-maze, mice freely explore each of three arms (labeled A, B, C). The behavioral output of this task is measured by the percentage of spontaneous alternations and percentage of re-entries into the same arm. A spontaneous alternation is defined as three consecutive entrances into novel arms without repeats (i.e. ABC, CBA). A re-entry is defined as entering the same arm twice in a row (i.e. AA, BB, CC). For chronic activation of CaMKII-specific Gq pathway, there was a significant interaction between group and time point for the percentage of spontaneous alterations (Two-way ANOVA; Interaction: F (2, 60) = 6.655, p=0.0025; Timepoint: F (2, 60) = 12.65, p<0.0001; Group: F (1, 60) = 0.9491, p=0.3339)(Supplemental Figure 1H). *Post hoc* analyses revealed that there were significant pairwise differences between the 3 vs. 6 month CaMKII-hM3Dq (Tukey’s: p<0.0001) and 3 vs. 9 month CaMKII-hM3Dq groups (Tukey’s: p=0.0002)(Supplemental Figure 1H). This suggests that our manipulation of CaMKII-specific Gq pathways and aging are interacting to drive this specific effect in y-maze behavior. Additionally, there was no interaction between group and time point for the number of re-entries in the CaMKII-hM3Dq and control groups. However, there was an independent effect of time point in this behavioral metric, suggesting that aging alone is contributing to a change in y-maze behavior (Twoway ANOVA; Interaction: F (2, 60) = 1.893, p=0.1596; Timepoint: F (2, 60) = 3.878, p=0.0261; Group: F (1, 60) = 0.2529, p=0.6169)(Supplemental Figure 1I).

For Gq activation of GFAP+ astrocytes, we observed no significant interaction between group and time point for percentage of spontaneous alternations or number of re-entries. However, there were main effects of time point and group for re-entries, suggesting that aging and our manipulation are contributing to a behavioral change in the y-maze (Two-way ANOVA; [Spontaneous alternations; Interaction: F (2, 56) = 1.085, p=0.3448; Timepoint: F (2, 56) = 3.015, p=0.0571; Group: F (1, 56) = 0.5859, p=0.4472][Re-entries; Interaction: F (2, 57) = 0.8932, p=0.4150; Timepoint: F (2, 57) = 3.183, p=0.0489; Group: F (1, 57) = 4.614, p=0.0360](Supplemental Figure 2H-I). Overall, CaMKII-Gq and GFAP-Gq activation compounded with aging had differential effects on y-maze behaviors.

### 3.8) CaMKII-hM3Dq activation changes the number of microglia in vHPC, but not the number of astrocytes. GFAP-hM3Dq activation does not change the number of microglia or astrocytes in vHPC

As glial cells play a major role in dysfunctional circuit activity in the brain (Liddelow et. al., 2020), we next quantified the cellular changes that our chronic astrocytic or neuronal manipulations induced. To accomplish this, we first quantified the total number of Iba1+ microglia in vCA1 across groups (Figure 6A-C; 7A-C). For CaMKII-hM3Dq and mCherry groups, there was no interaction between time point and group for the number of Iba-1+microglia in vCA1 of the HPC. However, there was an effect of our manipulation of the CaMKII-Gq pathway, suggesting that chronic activation of neurons in this region induces changes in microglial number (Two-way ANOVA; Interaction: F (2, 12) = 1.564, p=0.2492; Time point: F (2, 12) = 1.797, p=0.2077; Group: F (1, 12) = 10.86, p=0.0064)(Figure 6C). On the other hand, for the GFAP-hM3Dq and mCherry groups, there was no interaction between time point and group, nor were there individual effects of either factor on microglial number (Two-way ANOVA; Interaction: F (2, 12) = 0.5945, p=0.5673; Timepoint: F (2, 12) = 1.962, p=0.1831; Group: F (1, 12) = 0.3854, p=0.5463)(Figure 7C). Next, we quantified the total number of GFAP+ astrocytes in vHPC across groups. Across all CaMKII and GFAP groups, there were no significant interactions between time point and group for astrocytic number (Two-way ANOVA; [CaMKII-hM3Dq vs. mCherry: Interaction: F (2, 12) = 0.7052, p=0.5134; Timepoint: F (2, 12) = 3.220, p=0.0759; Group: F (1, 12) = 0.7784, p=0.3950][GFAP-hM3Dq vs. mCherry: Interaction: F (2, 12) = 0.2408, p=0.7897; Timepoint: F (2, 12) = 2.557, p=0.1189; Group: F (1, 12) = 2.672, p=0.1281])(Figure 6J, 7J). Overall, our CaMKII-hM3Dq manipulation induced significant changes in microglial number in vHPC, but did not affect astrocytic number. Additionally, our GFAP-hM3Dq manipulation did not impact microglial or astrocytic number.

**Figure 6.**
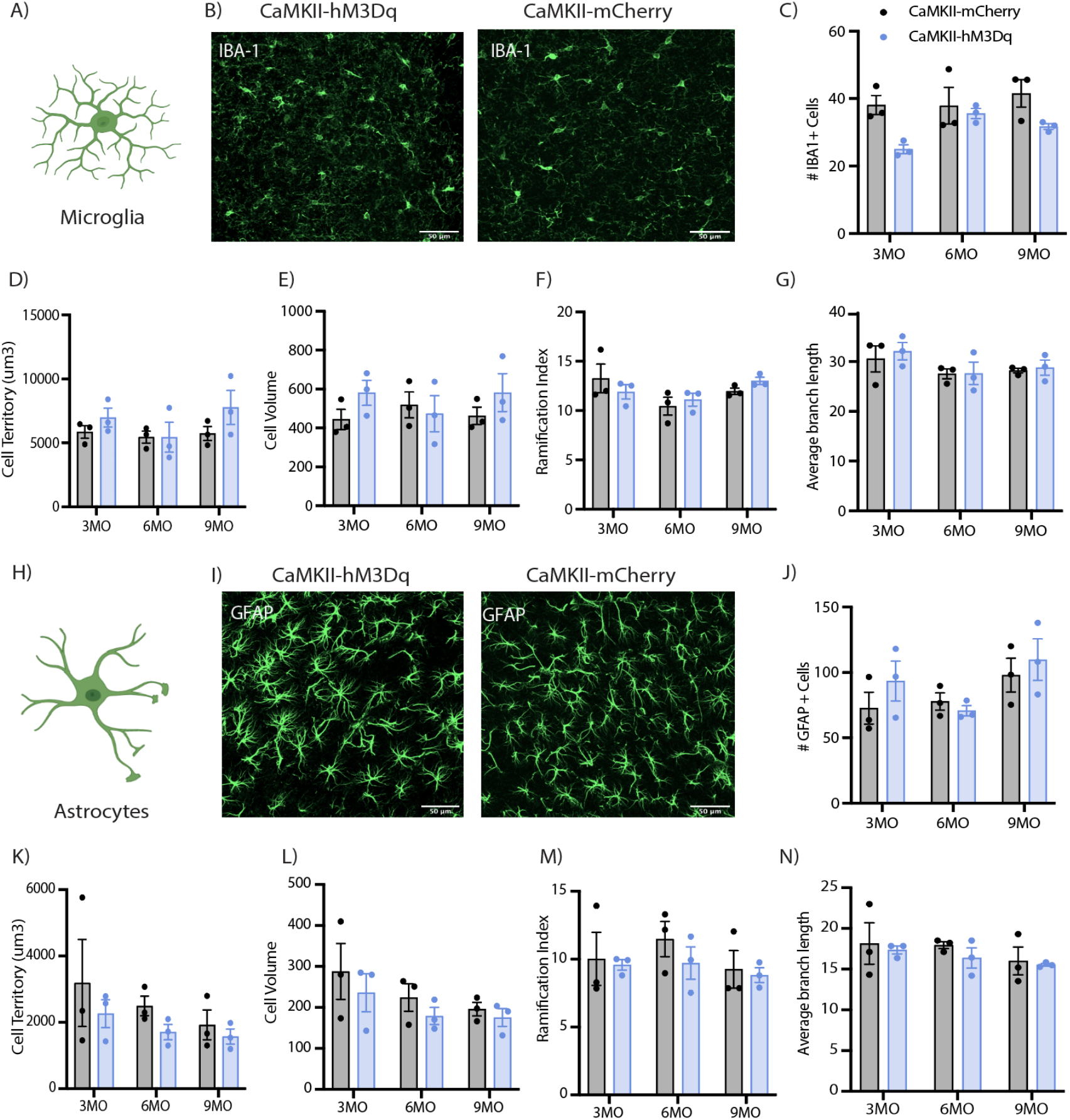
CaMKII-hM3Dq activation mildly impacted microglial cell number, but not astrocytic cell number or glial morphology. (A) Schematic of microglia morphology. (B) Representative images of Iba-1+ cells in GFAP-hM3Dq and GFAP-mCherry groups. (C) Number of Iba-1+ cells counted in 3, 6, and 9 month GFAP groups. (D-G) Quantification of Iba-1+ (D) cell territory volume, (E) cell volume, (F) ramification index, and (G) average branch length in 3, 6, and 9 month GFAP groups. (H) Schematic of astrocyte morphology. (I) Representative images of GFAP+ cells in GFAP-hM3Dq and GFAP-mCherry groups. (J) Number of GFAP+ cells counted in 3, 6, and 9 month GFAP-hM3Dq and GFAP-mCherry groups. (K-N) Quantification of GFAP+ (K) cell territory volume, (L) cell volume, (M) ramification index, and (N) average branch length in 3, 6, and 9 month GFAP groups. Cell count and morphology metrics were assessed with a two-way analysis of variance (ANOVA) with time point and group as factors. Tukey’s post hoc tests were performed where applicable. Error bars indicate SEM. p ≤ 0.05, **p ≤ 0.01, ***p ≤ 0.001, ****p ≤ 0.0001, ns = not significant. All groups: n= 3 mice x 18 tiles (ROI: vCA1) each were quantified for statistical analysis of astrocytic and microglial morphology per group.

### 3.9) GFAP-hM3Dq activation impacted microglial morphology, but CaMKII-hM3Dq did not

To further analyze the impact of our CaMKII or GFAP Gq pathway activation, we performed morphological analysis of microglia in vHPC. For CaMKII-hM3Dq and mCherry groups, there was no significant interaction between time point and group for microglial cell territory volume, cell volume, ramification index or average branch length (Two-way ANOVA; [Cell territory volume: Interaction: F (2, 12) = 0.7040, p=0.5139; Timepoint: F (2, 12) = 1.240, p=0.3239; Group: F (1, 12) = 2.244, p=0.1600][Cell volume: Interaction: F (2, 12) = 0.9601, p=0.4104; Timepoint: F (2, 12) = 0.06630, p=0.9362; Group: F (1, 12) = 1.423, p=0.2560][Ramification index: Interaction: F (2, 12) = 1.217, p=0.3303; Timepoint: F (2, 12) = 2.981, p=0.0889; Group: F (1, 12) = 0.02767, p=0.8707][Avg branch length: Interaction: F (2, 12) = 0.08429, p=0.9197; Timepoint: F (2, 12) = 2.464, p=0.1269; Group: F (1, 12) = 0.2552, p=0.6226])(Figure 6D-G). Additionally, for microglia there were no interactions between time point and group for the number of endpoints, minimum branch length, maximum branch length or number of branch points, but there was an effect of aging for the number of branch points and endpoints (Two-way ANOVA; [Num Endpoints: Interaction: F (2, 12) = 0.1217, p=0.8865; Timepoint: F (2, 12) = 4.365, p=0.0376; Group: F (1, 12) = 0.3498, p=0.5652][Min branch length: Interaction: F (2, 12) =0.4630, p=0.6402; Timepoint: F (2, 12) = 0.6621, p=0.5336; Group: F (1, 12) = 0.1133, p=0.7423][Max branch length: Interaction: F (2, 12) = 0.3205, p=0.7318; Timepoint: F (2, 12) = 2.398, p=0.1330; Group: F (1, 12) = 0.2729, p=0.6109][Num branch points: Interaction: F (2, 12) = 0.1286, p=0.8805; Timepoint: F (2, 12) = 4.621, p=0.0325; Group: F (1, 12) = 0.4333, p=0.5228])(Supplemental Figure 4K-N). Overall, Gq manipulation of CaMKII+ neurons did not have an effect on microglial morphology and only aging had an impact.

For GFAP-hM3Dq and mCherry groups, there was a significant interaction between group and time point for microglial cell territory volume (Two-way ANOVA; Interaction: F (2, 12) = 8.377, p=0.0053; Timepoint: F (2, 12) = 5.679, p=0.0184; Group: F (1, 12) = 0.4626, p=0.5093)(Figure 7D). Further analysis revealed that there were pairwise differences between the 3 vs. 6 month GFAP-hM3Dq and mCherry (Tukey’s: p=0.0258), and 3 vs. 9 month GFAP-hM3Dq and mCherry groups (Tukey’s: p=0.0085), suggesting that our manipulation and aging impacted morphological changes (Figure 7D). For microglial cell volume, there was only an effect of aging (Two-way ANOVA; Interaction: F (2, 12) = 2.353, p=0.1373; Timepoint: F (2, 12) = 5.674, p=0.0184; Group: F (1, 12) = 0.2007, p=0.6622)(Figure 7E). Further, there was a significant interaction between time point and group for microglial ramification index, suggesting that both aging and our manipulation impacted this morphological change (Two-way ANOVA; Interaction: F (2, 12) = 4.650, p=0.0320; Timepoint: F (2, 12) = 1.401, p=0.2838; Group: F (1, 12) = 1.751, p=0.2104)(Figure 7F). For average branch length, there was no significant interaction between time point and group (Two-way ANOVA; Interaction: F (2, 12) = 0.8177, p=0.4646; Timepoint: F (2, 12) = 2.600, p=0.1154; Group: F (1, 12) = 0.01994, p=0.8901)(Figure 7G). Additionally, there was no interaction between time point and group for the number of endpoints, minimum branch length, maximum branch length or number of branch points of microglia in the GFAP-hM3Dq and mCherry groups. There was only individual effects of time point for the minimum branch length (Two-way ANOVA; [Num endpoints: Interaction: F (2, 12) = 1.727, =0.2192; Timepoint: F (2, 12) = 1.292, p=0.3104; Group: F (1, 12) = 1.370, p=0.2645][Min branch length: Interaction: F (2, 12) = 1.935, p=0.1869; Timepoint: F (2, 12) = 0.04348, p=0.9576; Group: F (1, 12) = 1.729, p=0.2131][Max branch length: Interaction: F (2, 12) = 1.221, p=0.3291; Timepoint: F (2, 12) = 1.646, p=0.2335; Group: F (1, 12) = 0.1075, p=0.7487][Num branch points: Interaction: F (2, 12) = 1.645, p=0.2337; Timepoint: F (2, 12) = 1.192, p=0.3372; Group: F (1, 12) = 1.279, p=0.2801])(Supplemental Figure 4O-R). Overall, Gq manipulation of GFAP-Gq induced changes in microglial morphology that interacted with aging.

**Figure 7.**
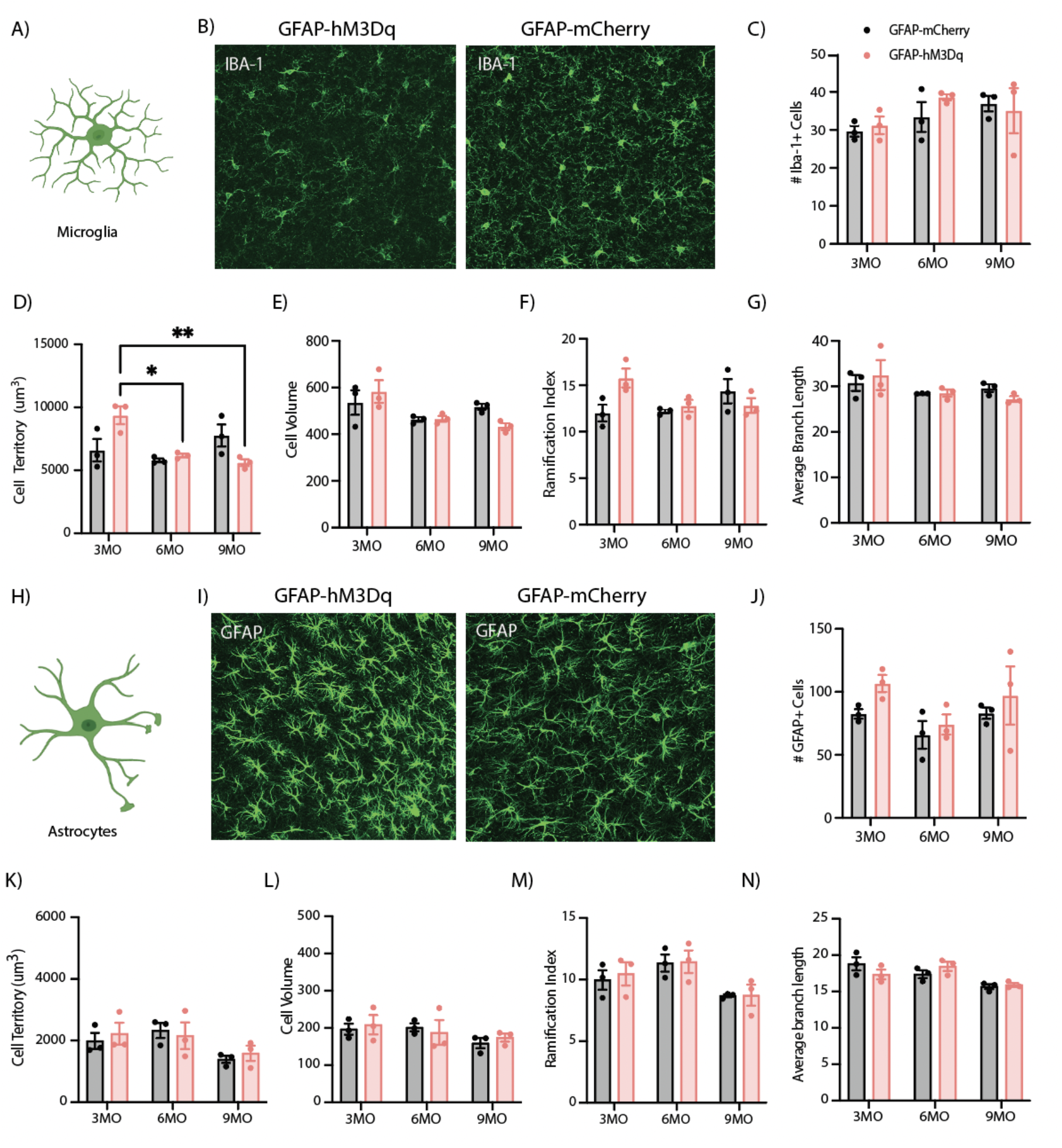
GFAP-hM3Dq activation did not affect glial cell number, but produced changes in microglial morphology. (A) Schematic of microglia morphology. (B) Representative single tile images of IBA-1+ cells in CaMKII-hM3Dq and CaMKII-mCherry groups. (C) Number of Iba-1+ cells counted in 3, 6, and 9 month CaMKII groups. (D-G) Quantification of Iba-1+ (D) cell territory volume, (E) cell volume, (F) ramification index, and (G) average branch length in 3, 6, and 9 month CaMKII groups. (H) Schematic of astrocyte morphology. (I) Representative images of GFAP+ cells in CaMKII-hM3Dq and CaMKII-mCherry groups. (J) Number of GFAP+ cells counted in 3, 6, and 9 month CaMKII groups. (K-N) Quantification of GFAP+ (K) cell territory volume, (L) cell volume, (M) ramification index, and (N) average branch length in 3, 6, and 9 month CaMKII groups. Cell count and morphology metrics were assessed with a two-way analysis of variance (ANOVA) with time point and group as factors. Tukey’s post hoc tests were performed where applicable. Error bars indicate SEM. p ≤ 0.05, **p ≤ 0.01, ***p ≤ 0.001, ****p ≤ 0.0001, ns = not significant. All groups: n= 3 mice x 18 tiles (ROI: vCA1) each were quantified for statistical analysis of astrocytic and microglial morphology per group.

### 3.10) Gq activation of GFAP+ or CaMKII+ cells in vHPC did not significantly affect astrocytic morphology, only aging had a mild impact in the GFAP group

To further analyze the impact of our CaMKII or GFAP Gq pathway activation on glial cells, we performed morphological analysis of astrocytes in vHPC. For CaMKII-hM3Dq and mCherry groups, there were no significant interactions between time point and group for astrocytic cell territory volume, total cell volume, ramification index, or average branch length (Two-way ANOVA; [Cell territory volume: Interaction: F (2, 12) = 0.1182, p=0.8896; Timepoint: F (2, 12) = 1.284, p=0.3125; Group: F (1, 12) = 1.850, p=0.1988][Cell volume: Interaction: F (2, 12) = 0.08956, p=0.9149; Timepoint: F (2, 12) = 2.126, p=0.1620; Group: F (1, 12) = 1.509, p=0.2428][Ramification index: Interaction: F (2, 12) = 0.1898, p=0.8296; Timepoint: F (2, 12) = 0.7775, p=0.4814; Group: F (1, 12) = 0.7513, p=0.4031][Avg branch length: Interaction: F (2, 12) = 0.08352, p=0.9204; Timepoint: F (2, 12) = 1.079, p=0.3707; Group: F (1, 12) = 0.6983, p=0.4197])(Figure 6K-N). Additionally, there was no interaction between time point and group for the number of endpoints, minimum branch length, maximum branch length or number of branch points for astrocytes in the CaMKII-hM3Dq and mCherry groups (Two-way ANOVA; [Num endpoints: Interaction: F (2, 12) = 0.4327, p=0.6585; Timepoint: F (2, 12) = 2.567, p=0.1180; Group: F (1, 12) = 1.718, p=0.2145][Min branch length: Interaction: F (2, 12) = 2.461, p=0.1272; Timepoint: F (2, 12) = 1.165, p=0.3449; Group: F (1, 12) = 0.03104, p=0.8631][Max branch length: Interaction: F (2, 12) = 0.1497, p=0.8626; Timepoint: F (2, 12) = 0.8872, p=0.4372; Group: F (1, 12) = 1.199, p=0.2950][Num branch points: Interaction: F (2, 12) = 0.4348, p=0.6572; Timepoint: F (2, 12) = 2.356, p=0.1370; Group: F (1, 12) = 1.539, p=0.2385])(Supplemental Figure 6K-N). Interestingly, there was an effect of group for minimum branch length, suggesting that our manipulation of the neuronal Gq pathway was affecting this characteristic of astrocytes independent of time point for this metric. Overall, Gq activation in CaMKII neurons only affects astrocytic minimum branch length, and no other metrics related to morphological changes.

For GFAP-hM3Dq and mCherry groups, there were no significant interactions between time point and group for astrocytic cell territory volume, total cell volume, ramification index, or average branch length. However, there was an effect of time point alone for both ramification index and average branch length, suggesting an impact of normal aging, but no effect of our Gq manipulation (Two-way ANOVA; [Cell territory volume: Interaction: F (2, 12) = 0.2921, p=0.7519; Timepoint: F (2, 12) = 3.777, p=0.0534; Group: F (1, 12) = 0.1344, p=0.7202][Cell volume: Interaction: F (2, 12) = 0.3062, p=0.7418; Timepoint: F (2, 12) = 1.796, p=0.2079; Group: F (1, 12) = 0.07025, p=0.7955][Ramification index: Interaction: F (2, 12) = 0.05066, p=0.9508; Timepoint: F (2, 12) = 6.080, p=0.0150; Group: F (1, 12) = 0.1177, p=0.7375][Avg branch length: Interaction: F (2, 12) = 2.351, p=0.1375; Timepoint: F (2, 12) = 9.220, p=0.0038; Group: F (1, 12) = 0.01233, p=0.9134])(Figure 7K-N). Additionally, there were no interactions between time point and group for the number of endpoints, minimum branch length, maximum branch length or the number of branch points for astrocytes in the GFAP-hM3Dq and mCherry groups (Two-way ANOVA; [Num endpoints: Interaction: F (2, 12) = 0.7147, p=0.5091; Timepoint: F (2, 12) = 4.391, p=0.0371; Group: F (1, 12) = 0.009931, p=0.9223][Min branch length: Interaction: F (2, 12) = 3.833, p=0.0516; Timepoint: F (2, 12) = 2.432, p=0.1298; Group: F (1, 12) = 0.1621, p=0.6943][Max branch length: Interaction: F (2, 12) = 1.563, p=0.2493; Timepoint: F (2, 12) = 7.779, p=0.0068; Group: F (1, 12) = 0.04654, p=0.8328][Num branchpoints: Interaction: F (2, 12) = 0.5939, p=0.5676; Timepoint: F (2, 12) = 4.401, p=0.0369; Group: F (1, 12) = 0.02290, p=0.8822])(Supplemental Figure 5O-R). Overall, unlike aging itself, Gq activation of the GFAP-Gq pathway does not affect any metrics of astrocytic morphological change.

## 4) Discussion

Our findings demonstrate that chronic activation of the Gq pathway in neurons and astrocytes differentially affects fear memory, social interaction, exploration, and anxiety-related behaviors, as well as markers of cellular stress. Together, our results add to burgeoning literature demonstrating that chemogenetic, optogenetic and pharmacological manipulation of neurons and astrocytes across brain regions induces behavioral enhancements or impairments (Nagai et. al., 2019; Chen et. al., 2016; Adamsky et. al., 2018; Martin-Fernandez et. al., 2017; Xiao et. al., 2020; Lei et. al., 2022; Shelkar et. al., 2021; Li et. al., 2020; Padilla-Coreano et. al., 2017; Deisseroth et. al., 2014; Jimenez et. al., 2018; Jennings et. al., 2013; Stuber et. al., 2012). Although these studies begin to address the role of these cell types in a variety of behaviors, there is a gap in our understanding of what prolonged or “chronic” manipulations do to network functioning as it relates to behavioral output. Our study aimed to address both of these gaps with chronic manipulation of the Gq pathway within vCA1.

vHPC is known to process contextual fear memory via communication with the PFC and amygdala, sending information about contextual representations and emotional valence. In this study, we find that CaMKII-hM3Dq activation in the vHPC decreased fear acquisition at 9 months and increased fear during extinction at 3 months. Recent literature has shown that stimulation of the vHPC at different frequencies can differentially impact freezing levels. For example, 20Hz stimulation decreases freezing levels via inhibition of basal amygdala (BA) excitatory responses (Graham et. al., 2021). Our stimulation of neurons at the 9 month time point may be impacting freezing levels during acquisition in a similar way via disruption of this monosynaptic fear pathway between vHPC and the amygdala. In this same study, lower frequency stimulation (i.e. 4Hz) showed a trend towards increased freezing during extinction sessions, providing a potential mechanistic understanding of our findings at 3 months (Graham et. al., 2021). An alternative explanation for our findings is derived from studies showing that lesions of the vHPC induce contextual freezing impairments by enhancing locomotor activity. If we disrupt this circuit, we may be enhancing locomotor activity and thus, decreasing freezing during acquisition without any real effects on memory processing itself (Richmond et. al., 1999). Another paper shows that vHPC double-projecting neurons to the PFC and amygdala are preferentially recruited during contextual fear memory acquisition (Kim & Cho, 2017). Our manipulation could be disrupting this contextual information transfer to both brain regions during acquisition and disturbing the synchronization that is necessary for updating the memory across extinction. For astrocytes, we find that GFAP-hM3Dq activation impacted CFC, recall and extinction at the 6 and 9 month time points. This is in line with recent literature supporting the importance of astrocytes in the successful acquisition and maintenance of memories in the hippocampus. For example, optogenetic activation of hippocampal astrocytes with channelrhodopsin (ChR2) during fear memory consolidation decreases fear and related anxiety behaviors in mice. Additionally, photostimulation of these cells rescues the decrease in distance traveled and time spent in the center of the open field that was produced by fear conditioning (Li et. al., 2020). Another study has shown that chemogenetic Gq activation or optogenetic stimulation of astrocytes in the dorsal hippocampus before or during fear conditioning enhances memory acquisition and subsequent recall via NMDA dependent LTP in dCA1 via D-serine release (Adamsky et. al., 2018). Our chronic Gq manipulation in the ventral hippocampus may be uniquely impacting memory in a manner that is mechanistically different from the described acute, dorsal hippocampal manipulations. These findings broadly suggest that different patterns of stimulation with opto- or chemogenetics in the hippocampus may induce differential effects on astrocytic signaling with neighboring neurons via unique release of gliotransmitters or be dependent on the brain region of interest.

Further, we find that both CaMKII-hM3Dq activation and aging exert a combinatory effect on anxiety-related behaviors across the open field and elevated zero maze. This is in line with previous work demonstrating that dentate gyrus (DG) stimulation of granule cells in the vHPC exerts anxiolytic effects, thus suppressing innate anxiety levels (Khierbeck et. al., 2013). Other work has shown that vCA1 is enriched with ‘anxiety cells’ that are activated by anxiogenic environments via projections to the lateral hypothalamus (LH) and are necessary for avoidance behaviors (Jimenez et. al., 2018). In addition, direct input from vHPC to the prefrontal cortex (PFC) is necessary for anxiety-related neural activity and inhibition of this projection decreases anxiety behaviors (Padilla-Coreano et. al., 2016; Ciocchi et. al., 2015). Together, this work allows us to understand how activation of neurons in this anxiety ‘hub’ through our manipulation could produce changes in anxiety. On the other hand, GFAP-hM3Dq activation decreased anxiety-related behaviors at the 3 month time point in the open field, but only showed impacts of aging in the zero maze. In light of recent literature, astrocytic activation with ChR2 increases calcium signaling intracellularly and reduces anxiety-like behaviors, thus suggesting that astrocytes play a role in overcoming or reducing anxiety levels (Cho et. al., 2022). Consistent with our results at the 3 month time point, optogenetic stimulation of astrocytes increases speed, distance and time spent in the center of the open field (Cho et. al., 2022). Another study showed that there is an interaction between anxiety and fear behaviors with astrocytic activation in the dorsal hippocampus, with activation decreasing anxiety-related behaviors in mice post-fear conditioning (Li et. al., 2020). In the context of Khierbeck et. al., 2013 discussed above, we believe that astrocytic stimulation and release of adenosine triphosphate (ATP) increases excitatory synaptic transmission in the ventral DG granule cells, together contributing to a suppression of innate anxiety (Cho et. al., 2022).

In the present study, we observed a decrease in social engagement in the CaMKII-hM3Dq mice at the 6 month time point that received chronic activation. GFAP-hM3Dq mice displayed only effects of aging on locomotion and social engagement. The findings of our neuronal activation are consistent with previous literature showing that vCA1 pyramidal cells projecting to the PFC to regulate mouse social behaviors (Sun et. al., 2020), as well as chemogenetic excitation of vCA1-PFC projecting cells impairing social memory (Philips et. al., 2019). Gq activation within the hippocampus may modulate neuronal outputs to these brain regions and impact social behaviors. Interestingly, novel environment exploration in the y-maze was impacted by a combination of chronic Gq activation and aging in both neuronal and astrocytic groups. Relatedly, recent findings suggest that stimulation of hippocampal astrocytes increases exploratory behaviors in anxiety-related tasks, making it difficult to fully disentangle a drive to explore a novel environment from innate anxiety levels (Cho et. al., 2022).

Glial cells, such as microglia and astrocytes exhibit unique changes morphologically, transcriptionally and functionally with aging, disease and cellular stress. In the dysfunctional human brain, glial cells become more vulnerable and may fail in their roles of neuronal protection, synaptic regulation and maintenance of homeostasis, gliotransmission and blood-brain barrier support. In the present study, we observe that CaMKII-hM3Dq mice have changes in microglial, but not astrocytic cell number in vHPC, while GFAP-hM3Dq mice showed no differences in glial cell number. In CaMKII- and GFAP-hM3Dq groups, we did not observe any changes in glial cell number with aging. This is consistent with literature showing that astrocytic staining does not show differences in the CA1 of aging mice (Grosche et. al., 2013). For microglia, studies have shown evidence that Iba1+ cell numbers in the hippocampus of old and young rats did not differ with aging (VanGuilder et. al., 2011). We speculate that changes in the number of microglia in vHPC from our neuronal manipulation may be due to efforts of the brain to reduce excitotoxicity induced by Gq activation over time.

In terms of morphological changes, CaMKII-hM3Dq activation did not have an impact, but aging affected the number of branch points and endpoints of microglia. GFAP-hM3Dq activation and aging combined to impact microglial cell volume, ramification index and minimum branch length. Studies have shown that older mice have a shortening and decrease in the complexity of microglial processes and branching structure (Hefendehl et. al., 2014). For astrocytes, CaMKII-hM3Dq activation minimally affected minimum branch length, and there was no effect of aging. Finally, GFAP-hM3Dq activation did not impact astrocytic characteristics, but aging had an effect on ramification index and the average branch lengths. Other work has provided evidence that astrocytes change morphology with aging (Ferrer et. al., 2017), modifying their GFAP+ surface area and volume in the hippocampus (Rodriguez et. al., 2014). A notable limitation of the quantification of glial cell number and morphology lies in the chosen cellular markers that were stained for each cell type. For example, GFAP expression does not allow for detection of fine astrocytic processes, providing conflicting results for the changes in area and complexity of these cells in the hippocampus with aging (Rodriguez et. al., 2014; Cerbai et. al., 2012).

Surprisingly, chronic manipulation of the Gq pathway in neurons and astrocytes may abolish some age-dependent changes in task performance that we observe in our control groups. For example, we observe an age-related increase in freezing behavior during CFC in our CaMKII-mCherry group that is abolished by our CaMKII-hM3Dq manipulation at the 9 month time point. Our manipulation may be working against these age-dependent changes by maintaining heightened levels of hippocampal activity, increased plasticity or activity-dependent increases in neurogenesis. Future research will be necessary to understand the underlying mechanisms of this age-related rescue with Gq activation and this may inform therapeutic interventions in an aging population that may rely on a mild increase in network activity within the hippocampus and/or neighboring brain regions.

On this note, our study provides evidence that cell-type specific targeting induces differential effects on behavioral outcome, which may help inform subsequent treatments for disorders of the brain. For instance, while deep brain stimulation (DBS) and transcranial magnetic stimulation (TMS) are often effective in the treatment of psychiatric disorders, the cell-types directly affected through each perturbation and their outcomes remain ripe for exploration. Along similar lines, recent literature has suggested that clinical DBS-like, high-frequency stimulation of human astrocytes promotes changes in gene expression that are relevant to extracellular matrix formation, likely aiding in synapse development and induction of neuronal plasticity (Jang et. al., 2019). Another example of this therapeutic potential is demonstrated by entorhinal cortex (EC) deep brain stimulation in 6 month old mice having the capacity to rescue memory deficits in a mouse model of neurodegeneration (Xia et. al., 2017). Further understanding of how these techniques may be non-discriminately targeting all cells within a brain region vs. specific cell-types to exert their effects on network functioning and improvement in symptomatology would improve their efficacy. Our work suggests that targeting astrocytes or neurons within the vHPC may be an effective means to differentially modulate fear, anxiety, exploratory, and social behaviors.

Chronic manipulation of Gq pathways in neurons and astrocytes across time has inherent limitations. DREADD-mediated cell activation has not been as thoroughly investigated with chronic use, as it has been in acute administration studies. As such, it is possible that across 9 months, the efficacy of the ligand binding decreases over time. However, and promisingly, we observe persistence in the receptor expression, as indicated by the robust expression of hM3Dq-mCherry across 3, 6 and 9 month time points in both cell types. Indeed, even if the DREADD-mediated manipulation was only robust until the 3 month time point, we are still inducing pronounced acute insults that impact behavior and cellular markers across all time points. Another important limitation of this study is the use of only male mice across all time points. In humans, although there is an increase in risk for women to develop neurodegenerative disease, males are more likely to develop severe cognitive and behavioral phenotypes. Specifically, males demonstrate increased aggression, shorter lifespan, increased severity of cognitive decline and earlier onset of Parkinson’s-associated dementia (Podcasy & Epperson, 2016). We chose to start with males and investigate the impact of both neuronal and astrocytic manipulation on behavior and cellular markers, but underscore the importance of work that will be needed to investigate the impact of chronic Gq pathway activation in females across cell types, together providing vital information on how male and female brains process network dysfunction differentially and offer insight into sex-specific therapeutic interventions for these devastating diseases and disorders that may result from this activity imbalance.

Finally, future research may investigate the *in vivo* physiological response using electrophysiology in vCA1 across time points as these cells are chronically activated. The impact of this would be two-fold: understanding how cellular activity changes of neurons in vCA1 change with manipulation across time, and confirming physiological response of these cells to the manipulation with a more concrete read-out than behavior. This would additionally allow for investigation of how cellular activity is related to the different behavioral changes (e.g. early changes in anxiety-related behaviors vs. later changes in fear memory). Overall, our data suggests that chronic manipulation of the Gq pathway in neurons and astrocytes across multiple time points impacts behavior. The results presented here provide valuable insights into the differential effects of chemogenetic manipulations of multiple cell types in a single brain region, as well as the specific effects of network disruption in the vHPC on cellular and behavioral phenotypes.

## Data Availability Statement

The data that support the findings of this study are available from the corresponding author upon reasonable request.

## Acknowledgements

This work was supported by a Ludwig Family Foundation grant, an NIH Early Independence Award (DP5 OD023106-01), an NIH Transformative R01 Award, a Young Investigator Grant from the Brain and Behavior Research Foundation, the McKnight Foundation Memory and Cognitive Disorders award, the Pew Scholars Program in the Biomedical Sciences, the Air Force Office of Scientific Research (FA9550-21-1-0310), the Center for Systems Neuroscience and Neurophotonics Center at Boston University.

---

**Supplemental Figure 1:**
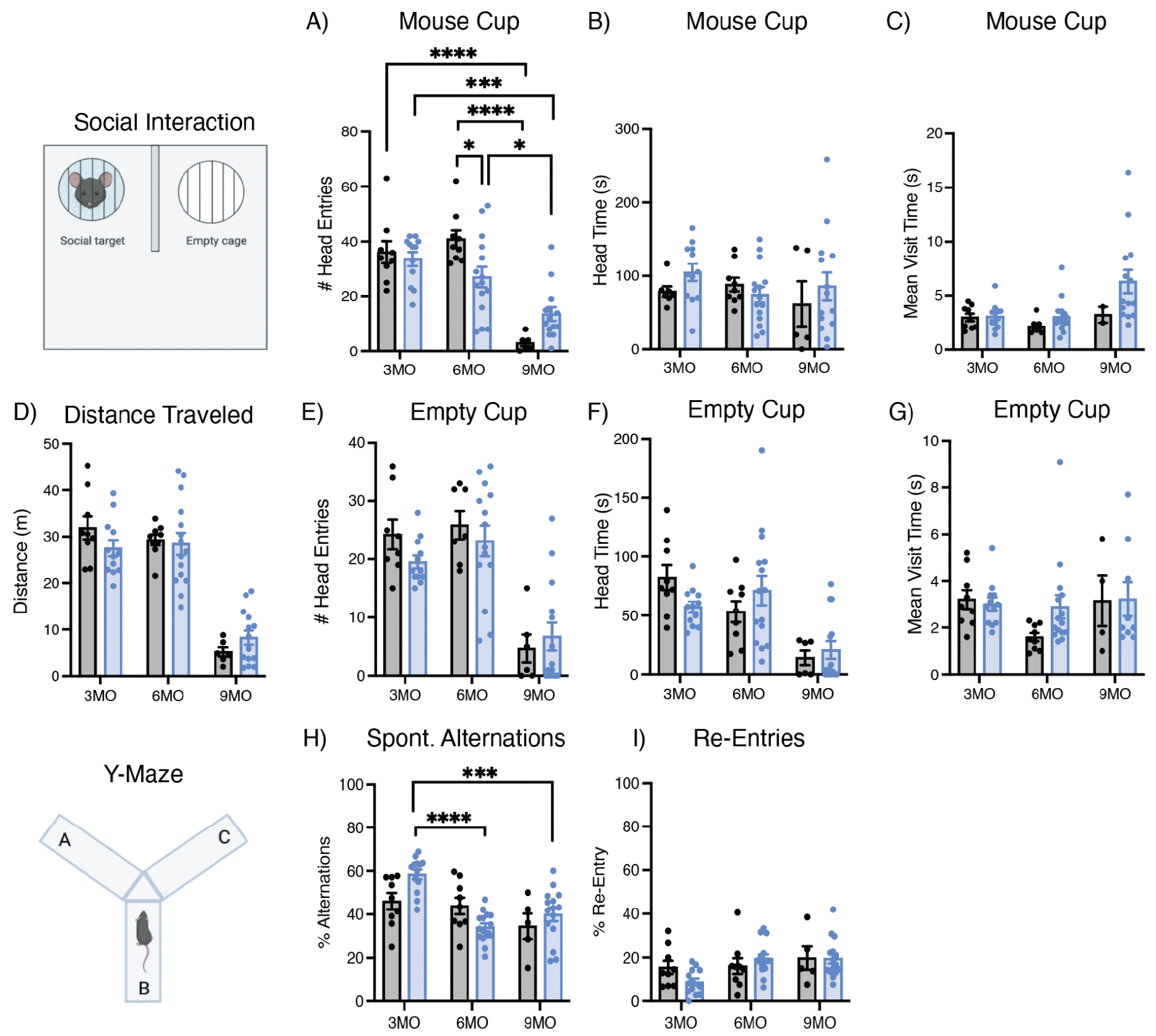
Chronic Gq activation of neurons in vCA1 decreases social engagement at the 6 month time point, and impacts y-maze behavior in combination with aging. (A-C) Social interaction mouse cup (e.g. social target) number of head entries (A), head time (B), and mean visit time (C) across the 3, 6 and 9 month CaMKII-mCherry and hM3Dq groups. (D) Total distance traveled across the entire testing chamber. (E-G) Social interaction empty cup (i.e. control target) number of head entries (E), head time (F), and mean visit time (G) across the 3, 6 and 9 month CaMKII-mCherry and hM3Dq groups. (H-I) Elevated y-maze percentage of spontaneous alternations (e.g. sequence of non-repeating arms; ABC, CBA) (H) and percentage of re-entries into the same arm (e.g. AA, BB, CC). Social interaction and y-maze metrics were assessed with a two-way analysis of variance (ANOVA) with time point and group as factors. Tukey’s post hoc tests were performed where applicable. Error bars indicate SEM. p ≤ 0.05, **p ≤ 0.01, ***p ≤ 0.001, ****p ≤ 0.0001, ns = not significant. For social interaction, 3 Month: hM3Dq (n=12), mCherry (n=7-9); 6 Month: hM3Dq (n=14-15), mCherry (n=8-9); 9 Month: hM3Dq (n=14), mCherry (n=3-6) after outlier removal. For y-maze, 3 Month: hM3Dq (n=13), mCherry (n=9); 6 Month: hM3Dq (n=15), mCherry (n=9); 9 Month: hM3Dq (n=15), mCherry (n=5) after outlier removal.

**Supplemental Figure 2:**
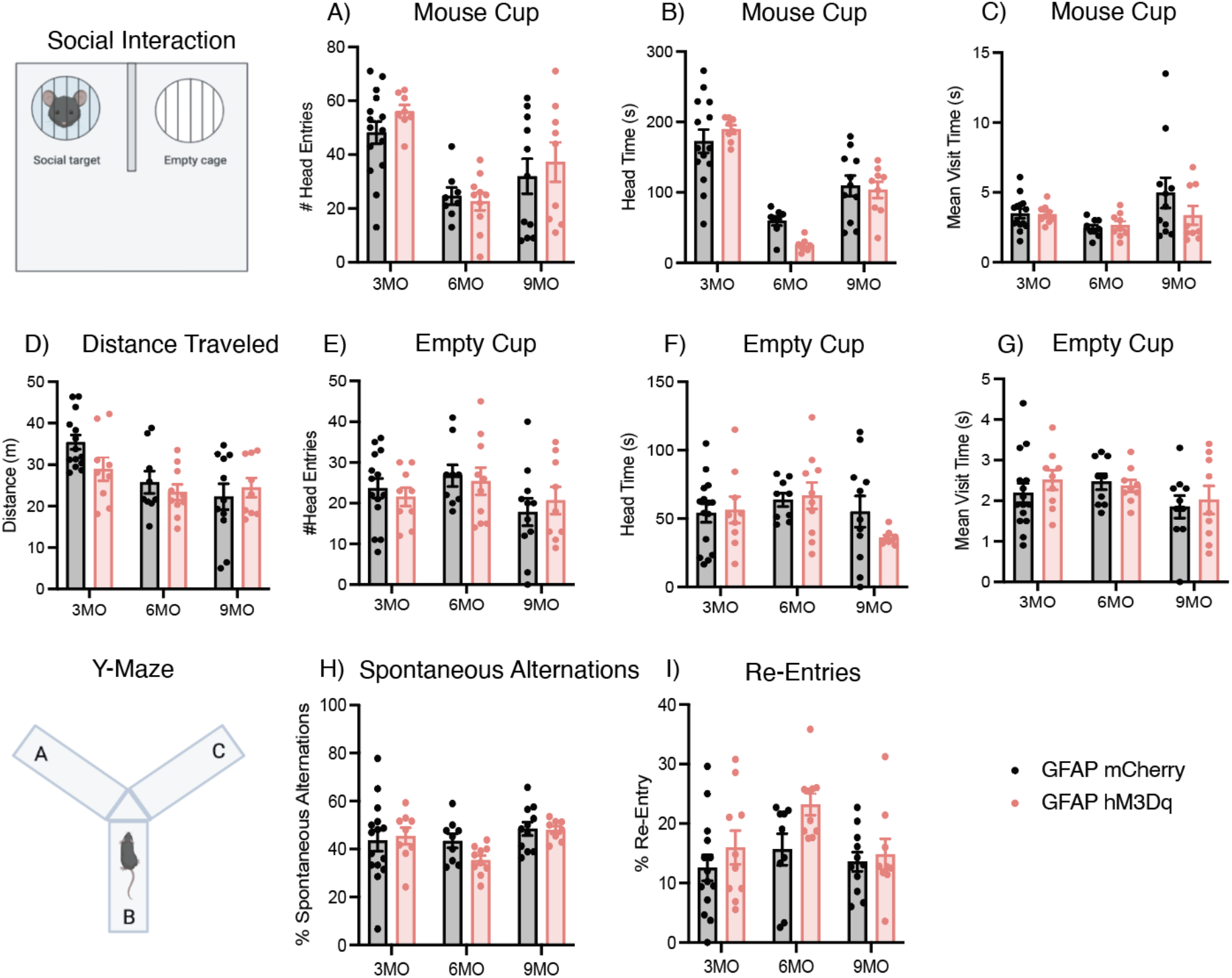
Chronic Gq activation of astrocytes in vCA1 does not impact social interaction, only aging has an effect. Y-maze behavior is impacted by chronic activation and aging. (A-C) Social interaction mouse cup (e.g. social target) number of head entries (A), head time (B), and mean visit time (C) across the 3, 6 and 9 month GFAP-mCherry and hM3Dq groups. (D) Total distance traveled across the entire testing chamber. (E-G) Social interaction empty cup (i.e. control target) number of head entries (E), head time (F), and mean visit time (G) across the 3, 6 and 9 month GFAP-mCherry and hM3Dq groups. (H-I) Elevated y-maze percentage of spontaneous alternations (e.g. sequence of non-repeating arms; ABC, CBA) (H) and percentage of re-entries into the same arm (e.g. AA, BB, CC). Social interaction and y-maze metrics were assessed with a two-way analysis of variance (ANOVA) with time point and group as factors. Tukey’s post hoc tests were performed where applicable. Error bars indicate SEM. p ≤ 0.05, **p ≤ 0.01, ***p ≤ 0.001, ****p ≤ 0.0001, ns = not significant. For social interaction, 3 Month: hM3Dq (n=8-9), mCherry (n=14); 6 Month: hM3Dq (n=9-10), mCherry (n=8-9); 9 Month: hM3Dq (n=8-9), mCherry (n=10-11) after outlier removal. For y-maze, 3 Month: hM3Dq (n=9-10), mCherry (n=14-15); 6 Month: hM3Dq (n=10), mCherry (n=9); 9 Month: hM3Dq (n=8-9), mCherry (n=11) after outlier removal.

**Supplemental Figure 3:**
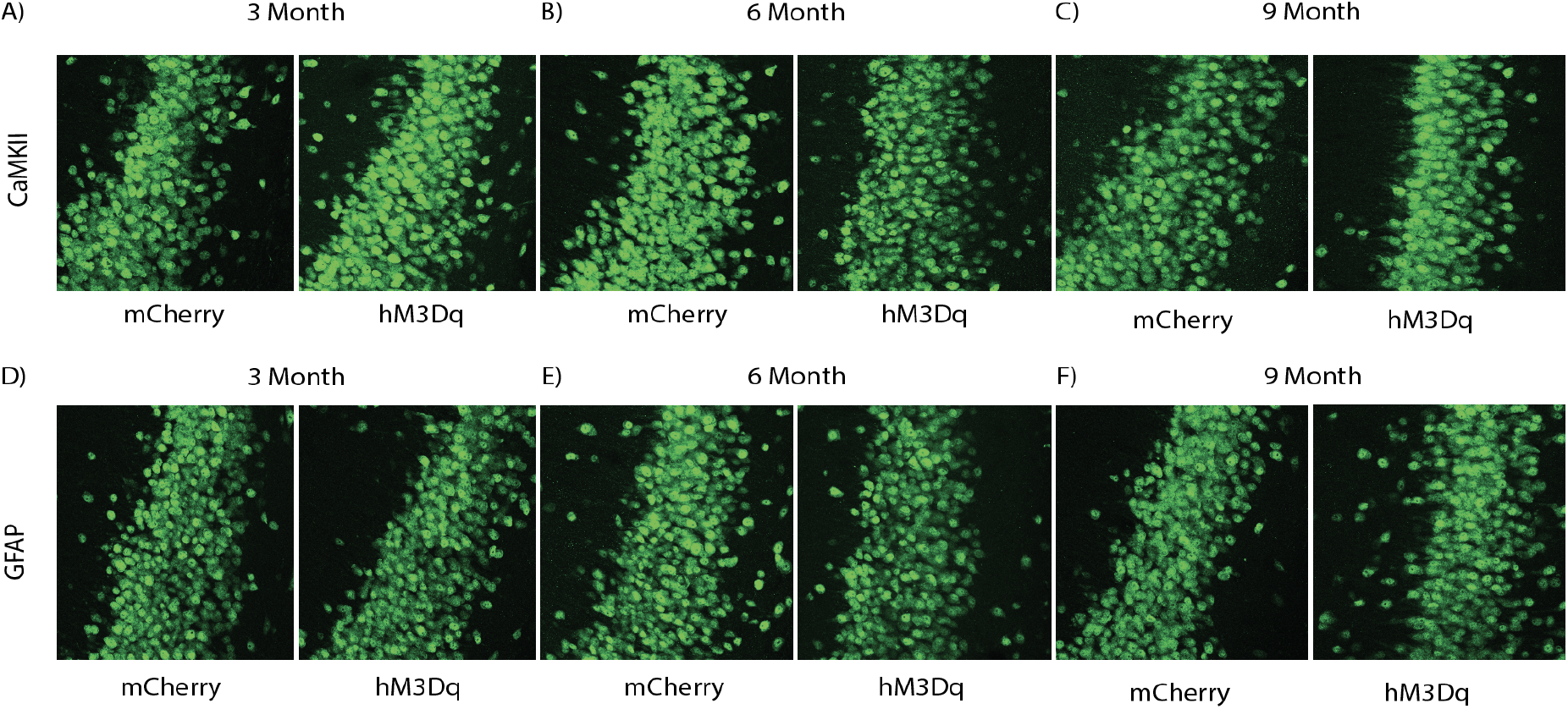
CaMKII-Gq pathway activation for 3, 6 or 9 months does not increase cell death in the vCA1 region of the hippocampus, but GFAP-Gq activation does in an aging-independent manner. (A-C) Representative hippocampal ventral CA1 NeuN (green) expression for CaMKII-mCherry and hM3Dq groups for the 3 month (A), 6 month (B) and 9 month (C) timepoints to quantify cell loss in the region of manipulation. (D-F) Representative hippocampal ventral CA1 NeuN expression for GFAP-mCherry and hM3Dq groups for the 3 month (D), 6 month (E) and 9 month (F) timepoints.

**Supplemental Figure 4:**
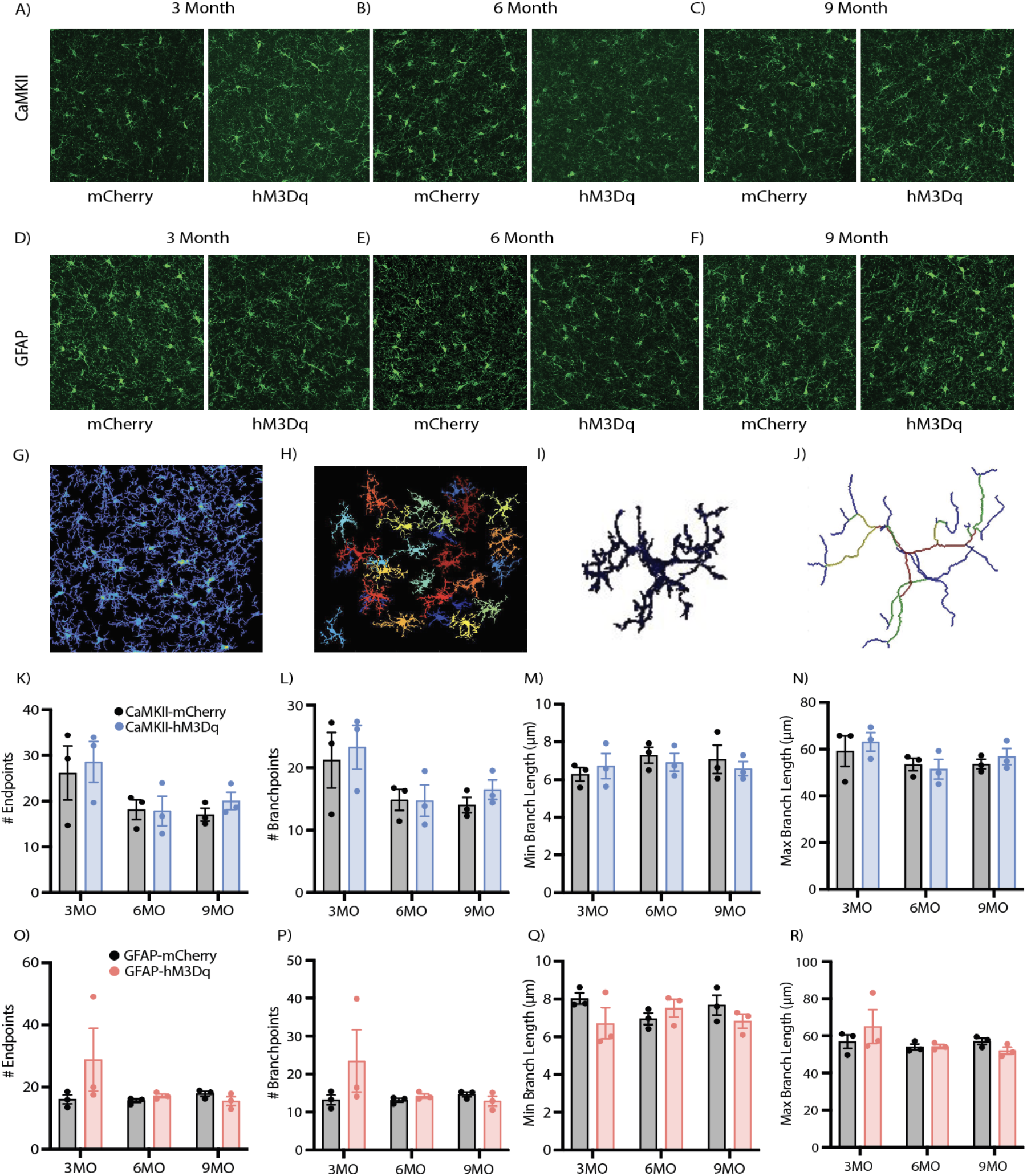
Microglial morphology in vCA1 is mildly impacted only by Gq activation of GFAP+ cells in vHPC. (A-C) Representative hippocampal ventral CA1 Iba-1 (green) expression for CaMKII-mCherry and hM3Dq groups for the 3 month (A), 6 month (B) and 9 month (C) timepoints to quantify cell number and morphological changes between groups and across time points. (D-F) Representative hippocampal ventral CA1 Iba-1 (green) expression for GFAP-mCherry and hM3Dq groups for the 3 month (D), 6 month (E) and 9 month (F) timepoints. (GJ) Representative image processing in 3DMorph; 2-dimensional (2D) threshold map generated from threshold and noise filtering parameters of a 3-dimensional (3D) z-stack image (G), isolated full cells generated by maximum and minimum cell size parameters (H), individual full cell (I), and skeletonized full cell used to generate outputs (e.g. territorial volume, cell size, ramification index) (J). (K-N) Quantification of remaining Iba-1+ morphological metrics; number of endpoints (K), number of branch points (L), minimum branch length (M) and maximum branch length (N) for the 3, 6 and 9 month CaMKII-mCherry and hM3Dq groups. (O-R) Quantification of remaining Iba-1+ morphological metrics; number of endpoints (O), number of branch points (P), minimum branch length (Q) and maximum branch length (R) for the 3, 6 and 9 month GFAP-mCherry and hM3Dq groups. Microglial morphology metrics were assessed with a two-way analysis of variance (ANOVA) with time point and group as factors. Tukey’s post hoc tests were performed where applicable. Error bars indicate SEM. p ≤ 0.05, **p ≤ 0.01, ***p ≤ 0.001, ****p ≤ 0.0001, ns = not significant. All groups: n= 3 mice x 18 tiles (ROI: vCA1) each were quantified for statistical analysis of astrocytic and microglial morphology per group.

**Supplemental Figure 5:**
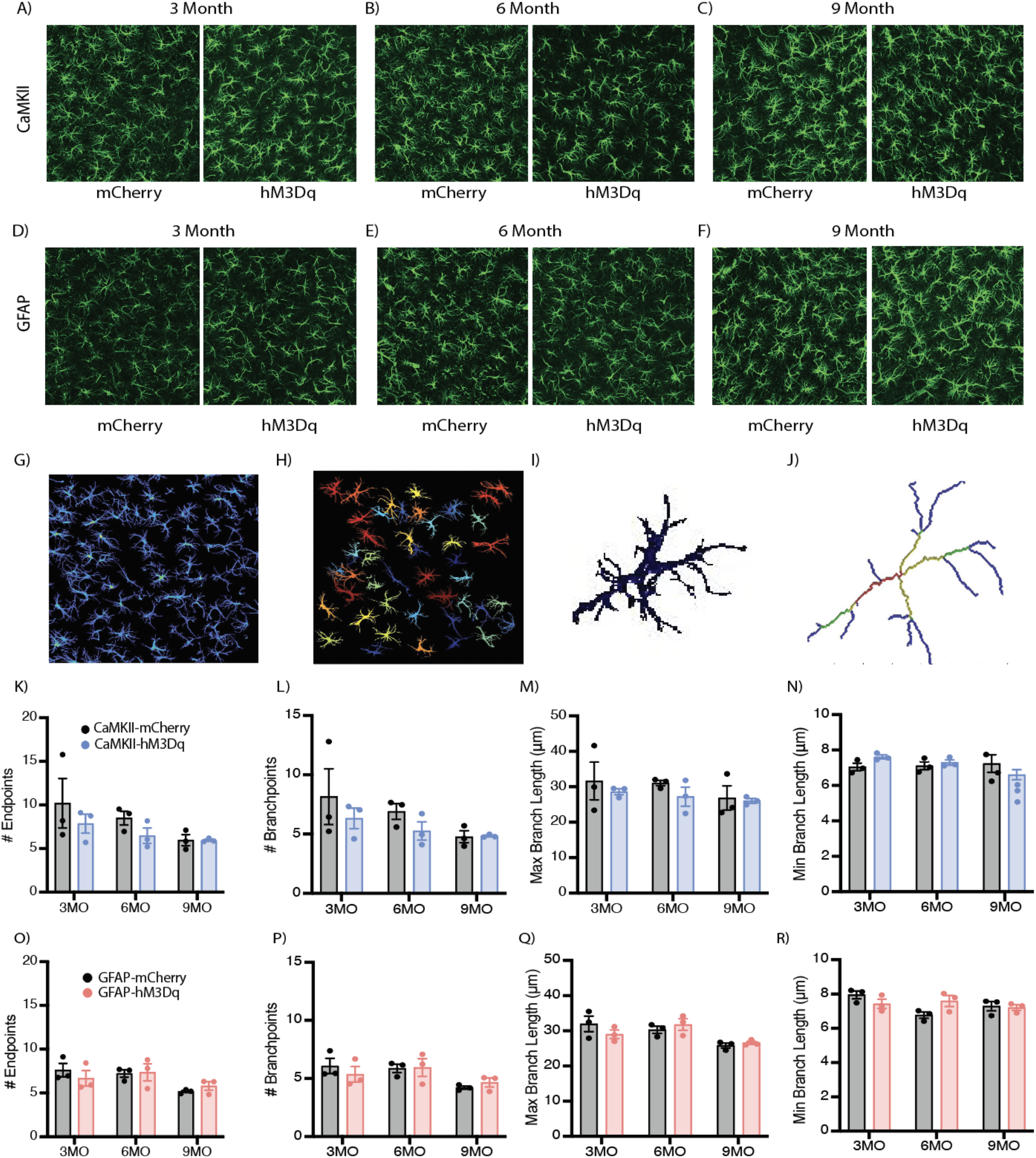
Astrocytic morphology in vCA1 showed no significant differences with Gq activation of CaMKII+ or GFAP+ cells in vHPC. (A-C) Representative hippocampal ventral CA1 GFAP (green) expression for CaMKII-mCherry and hM3Dq groups for the 3 month (A), 6 month (B) and 9 month (C) timepoints to quantify cell number and morphological changes between groups and across time points. (D-F) Representative hippocampal ventral CA1 GFAP (green) expression for GFAP-mCherry and hM3Dq groups for the 3 month (D), 6 month (E) and 9 month (F) timepoints. (G-J) Representative image processing in 3DMorph; 2-dimensional (2D) threshold map generated from threshold and noise filtering parameters of a 3-dimensional (3D) z-stack image (G), isolated full cells generated by maximum and minimum cell size parameters (H), individual full cell (I), and skeletonized full cell used to generate outputs (e.g. territorial volume, cell size, ramification index) (J). (K-N) Quantification of remaining GFAP+ cell morphological metrics; number of endpoints (K), number of branch points (L), minimum branch length (M) and maximum branch length (N) for the 3, 6 and 9 month CaMKII-mCherry and hM3Dq groups. (O-R) Quantification of remaining GFAP+ cell morphological metrics; number of endpoints (O), number of branch points (P), minimum branch length (Q) and maximum branch length (R) for the 3, 6 and 9 month GFAP-mCherry and hM3Dq groups. Astrocytic morphology metrics were assessed with a two-way analysis of variance (ANOVA) with time point and group as factors. Tukey’s post hoc tests were performed where applicable. Error bars indicate SEM. p ≤ 0.05, **p ≤ 0.01, ***p ≤ 0.001, ****p ≤ 0.0001, ns = not significant. All groups: n= 3 mice x 18 tiles (ROI: vCA1) each were quantified for statistical analysis of astrocytic and microglial morphology per group.

**Supplemental Figure 6.**
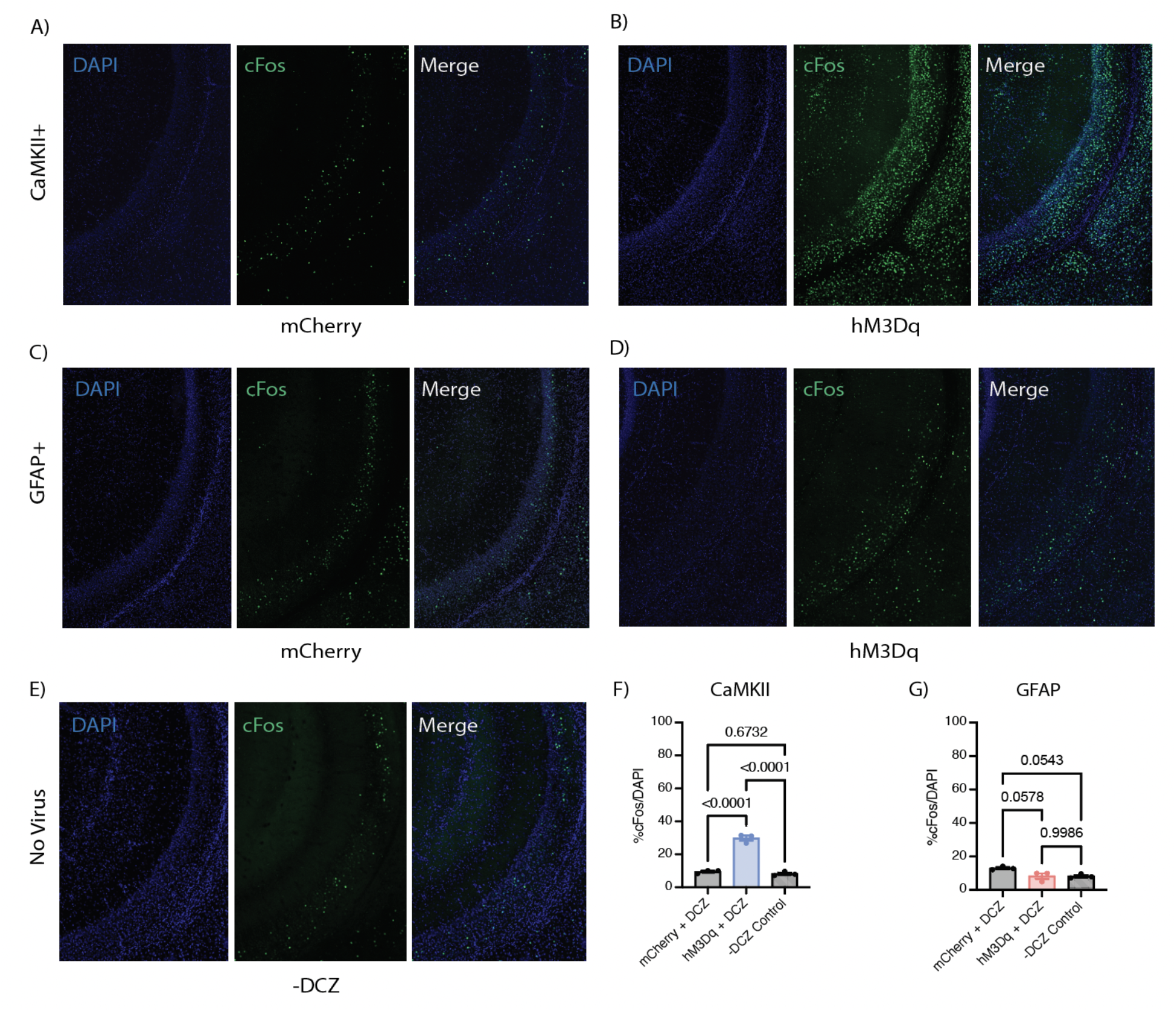
Gq activation with intraperitoneal (I.P.) administration of deschloroclozapine (DCZ) at 0.3mg/kg in GFAP-hM3Dq+ cells decreases vHPC cFos levels in the pyramidal cell layer, while activation of CaMKII-hM3Dq+ cells increases cFos levels. (A-B) Representative hippocampal ventral CA1 expression of DAPI and cFos in the CaMKII-mCherry (left) and CaMKII-hM3Dq (right) groups. (C-D) Representative hippocampal ventral CA1 expression of DAPI and cFos in the GFAP-mCherry (left) and GFAP-hM3Dq (right) groups. (E) Representative hippocampus vCA1 ‘baseline’ expression of DAPI and cFos in mice without virus or DCZ administration. (F-G) Quantification of % cFos/DAPI in the vCA1 pyramidal layer for (F) CaMKII-mCherry, CaMKII-hM3Dq and -DCZ groups and (G) GFAP-mCherry, GFAP-hM3Dq and -DCZ groups. All groups: n= 3 mice x 4 slices (ROI: vCA1 pyramidal cell layer) each were quantified for statistical analysis of cFos/DAPI counts per group. Error bars indicate SEM. For independent t-tests, p ≤ 0.05, **p ≤ 0.01, ***p ≤ 0.001, ****p ≤ 0.0001, ns = not significant.

**Supplemental Figure 7.**
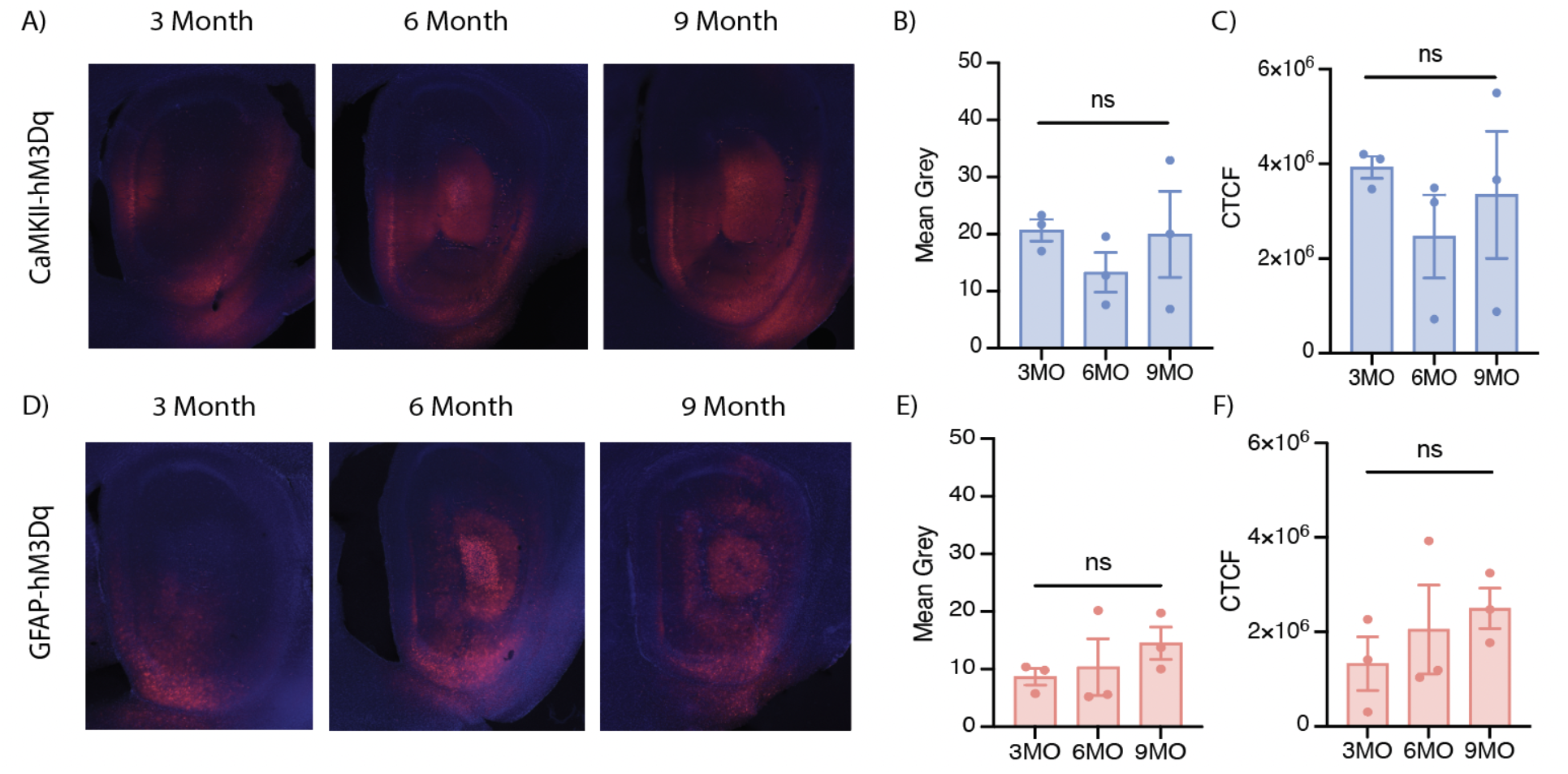
Gq receptor fluorescence is maintained across the 3, 6 and 9 month time points in both CaMKII and GFAP groups. (A-C) Representative ventral hippocampal Gq-mCherry fluorescence intensity for CaMKII-hM3Dq groups for the 3 month, 6 month, and 9 month time points to quantify expression levels using (B) Mean gray and (C) Corrected total cell fluorescence (CTCF). (D-F) Representative ventral hippocampal Gq-mCherry fluorescence intensity for GFAP-hM3Dq groups for the 3 month, 6 month, and 9 month time points to quantify expression levels using (E) Mean gray and (F) Corrected total cell fluorescence (CTCF). All groups: n= 3 mice x 3 vHPC slices each were quantified for statistical analysis using a one-way analysis of variance (ANOVA) for the mean gray or CTCF across time points. Tukey’s post hoc tests were performed where applicable. Error bars indicate SEM and p ≤ 0.05, **p ≤ 0.01, ***p ≤ 0.001, ****p ≤ 0.0001, ns = not significant.

